# Integrative assessment of the transcriptome and virome of the poultry red mite *Dermanyssus gallinae*

**DOI:** 10.1101/2022.04.21.489001

**Authors:** José M Ribeiro, David Hartmann, Pavla Bartošová-Sojková, Humberto Debat, Martin Moos, Petr Šimek, Jiří Fara, Martin Palus, Matěj Kučera, Ondřej Hajdušek, Daniel Sojka, Petr Kopáček, Jan Perner

## Abstract

*Dermanyssus gallinae* is a blood-feeding mite that parasitises on wild birds and farmed poultry. The *D. gallinae* mite has a short life cycle of fewer than two weeks from the egg to an egg-laying female. The remarkably swift processing of blood, together with the capacity to blood-feed in most developmental stages, makes this mite a highly debilitating pest. We have constructed developmental stage-specific transcriptomes, through Illumina RNA-seq, to mine the repertoire of protein-encoding mRNA transcripts, products of which participate in key processes that ensure the success of blood digestion, rapid ontogeny, and immunity. As a result of high reproductive capacity, the prevalence of *D. gallinae* in egg-producing poultry farms globally causes significant economic losses. Acaricides that are used to limit the reproduction of *D. gallinae* mites target cys-loop ion channels, which are widely shared across the phylogeny of invertebrates. To catalogue a comprehensive list of potential invertebrate-specific ion channels, we have constructed and analysed an additional RNA-seq library of *D. gallinae* micro-dissected midguts, a tissue with direct exposure to host blood and potential anti-parasitics. We phylogenetically defined groups of cys-loop proteins and probed their sensitivity to selected acaricides. Ultimately, we have catalogued all assembled transcripts and their expression values in a hyper-linked excel sheet with available sequences of individual contigs. The transcriptomic data were complemented by mass-spectrometry (MS)-based metabolite identification and by viability assays using selected inhibitors applied either by microinjection or through artificial membrane feeding. Additionally, we have described the RNA-virome of *D. gallinae* and identified a novel virus dubbed Red Mite Quaranjavirus 1.

## Introduction

The *Dermanyssus* mites are blood-feeding ectoparasites of birds ^1^. The poultry red mite (*D. gallinae*) is a global pest in layer houses for both domestic and commercial intensive egg production ^2,3,4^, which are part of an important and ever-increasing global market ^5^. *D. gallinae* mites have a very short life cycle, going from juveniles to reproducable adult stages within one week. The necessity to blood feed for most developmental stages, and the swift reproductive dynamics makes *D. gallinae* a highly irritating and troublesome pest. Within the first hour of the mite’s infestation, lymphocytic infiltration occurs and is a constant pathological feature, concomitant with necrosis of feather follicles, which is later followed by hyperkeratosis, parakeratosis, and acanthosis ^6^.

Despite general knowledge of the impact of mite infestation, our understanding of molecular processes enabling swift and repetitive blood-feeding, blood digestion, development and reproduction remains scarce. To bridge this knowledge gap, we report here on the mite’s transcriptomic blue-prints integral for each developmental stage. Special attention was paid to processes inherent for successful blood-feeders, including proteolytic blood digestion, haem and iron biology, feeding status signaling, vitellogenesis, and immunity. The divergence of cys-loop ligand-gated ion channels, the sensitive acaricide targets, is also described. The RNA-seq data are also complemented by viability assays, whereby small-molecule inhibitors are introduced to the mite by *ex vivo* membrane feeding or microinjection into its haemocoel.

During blood-feeding, *D. gallinae* mites can transmit several significant animal pathogens to their hosts ^7^, including some that are zoonotic ^8^. Although a large number of viruses and bacteria have been found associated with *D. gallinae*, its capacity to act as a vector or reservoir was experimentally supported for a few pathogens ^9-11^. This is especially alarming for transmission of *Salmonella* spp.^11^, causing the egg-associated salmonellosis and fowl typhoid disease ^12^, as well as of avian influenza A virus ^13^. *D. gallinae* has also been associated with several other bacterial and viral species, with proof awaiting experimental confirmation of its vectorial capacity ^2,11,14,15^. To reflect on the RNA virome currently present in *D. gallinae* mites, we filtered out individual viral sequences. Expression values of some viral sequences indicate active infection, which warrants further research into confirmation of vector/reservoir relevancy of *D. gallinae* mites.

## Results and Discussion

### *D. gallinae* transcriptome assembly and development of the BLASTable dataset

Transcriptome composition of four developmental stages of *D. gallinae* and micro-dissected midguts were investigated in this work by *de novo* assembly of Illumina RNA-seq reads. Specifically, we have constructed RNA-seq libraries of whole bodies of unfed protonymphs (naive pre-blood-feeding stage), fed protonymphs, fed deutonymphs, fed adults, and midguts dissected from fed adults (**Fig. 1A**). For each library, > 54 M reads with Phred Scores of Q30 ≥ 93.90 % were obtained (**Fig. 1B**). These were assembled into 85,117 contigs (Supplementary Data S1). Following their annotation, bacterial and chicken contaminants were removed. As *D. gallinae* mites were fed chicken blood, consisting of nucleated white and red cells, a majority of hen-specific reads (79%) were found, as expected, in the transcriptome of the midgut of adult *D. gallinae* mites **(Fig 1C**). Contigs were also found matching viruses, accruing 2.2% of the total reads (**Fig. 1C**). A total of 18,101 *D. gallinae*-specific contigs were deposited at the NCBI server as BioProject PRJNA597301 and Transcriptome Shotgun Assembly (TSA) GIFZ00000000 and are accessible through NCBI BLAST of the TSAdatabase. The contigs are also listed in a hyper-linked Excel sheet (see **Data availability**) with available encoded predicted protein characteristics, annotations of assembled contigs according to a particular database, differential expression statistics, and predicted cellular process involvement. To assess the completeness of the transcriptome, we ran a BUSCO analysis of the proteome, the result of which exhibited a yield of 91.8% complete BUSCOs.

**Figure 1.**
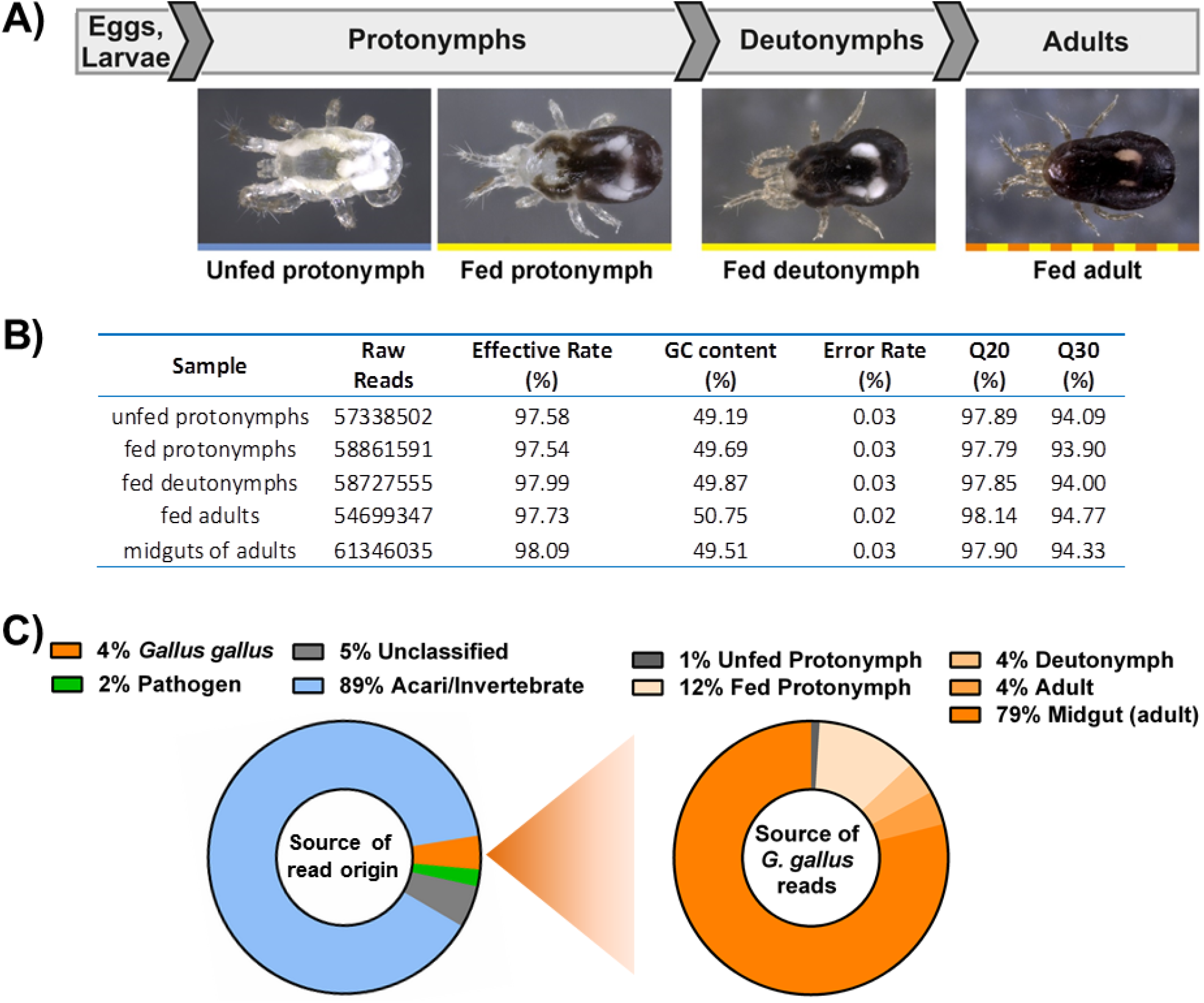
Schematic illustration of the experimental design. **A)** Mites were collected and sorted according to their developmental stages, and protonymphs also according to their feeding status. **B)** Salient information on read counts and quality scores achieved by Illumina sequencing. **C)** Attributed origin of the reads, from all libraries, according to the matched organism (left). The library origin of the chicken(*Gallus gallus*)-derived contigs is shown on the right.

### Core transcriptome, ontogenic idiosyncrasies, and blood-feeding associated coding transcripts

Larvae of *D. gallinae* do not feed on host blood, but the nymphal stages do. They feed once only, in order to develop into subsequent stages, i.e. protonymphs into deutonymphs, and deutonymphs into adults. Adult females then feed blood repeatedly, with each blood-feeding being followed by oviposition. Here, we used two comparative models: (*i*) comparing developmental stage-specific transcriptomes and (*ii*) comparing unfed and blood-fed mites. As for (*i*), the generated heat map clearly indicates differences in transcriptomes of juvenile and adult stages (**Fig. 2A**). We have pinned down similarities in individual expression patterns of transcripts between libraries of juvenile stages and the distinct expression profile of adults (**Fig. 2B**). We have identified 7,252 coding sequences (65.2%) shared across developmental stages (**Fig. 2B;** Supplementary Data S2) and these represent the transcriptomic core of *D. gallinae* mites. Indeed, immature stages (protonymphs and deutonymphs) share more transcripts with each other than any of the immature stages with adults (**Fig. 2B**). Each stage expresses 4 – 5.5% of unique idiosyncratic transcripts (**Fig. 2B;** Supplementary Table S2). Libraries of adult mites were characterised by highly abundant transcripts encoding proteins that participate in nuclear regulation and nutrient storage, both of which are essential for the reproduction of mites (**Fig. 2C;** Supplementary Data S3). As for (*ii*) we have compared transcriptomes of fed-vs unfed-protonymphs, and assessed their midgut-specific expression (**Fig. 2D**), in order to identify the blood feeding/digestion-associated set of encoded proteins. To identify individual proteins integral to blood-feeding of mites, we have filtered transcripts that were at least 16 times more abundant in fed protonymphs than in unfed protonymphs. These transcripts were further enriched for those having at least an FPKM value of 1 in the midgut transcriptomic library. In this way, we have identified 264 transcripts, which we propose encode blood-feeding-associated proteins (**Fig. 2E;** Supplementary Data S4). Of these, 143 transcripts (54 %) were annotated with a clear protein class assignment. The gene ontology indicates enrichment in signaling, nuclear regulation of expression, and proteostasis, with the extracellular matrix, signaling, and proteostasis comprising the most abundant transcripts (**Fig. 2F**).

**Figure 2.**
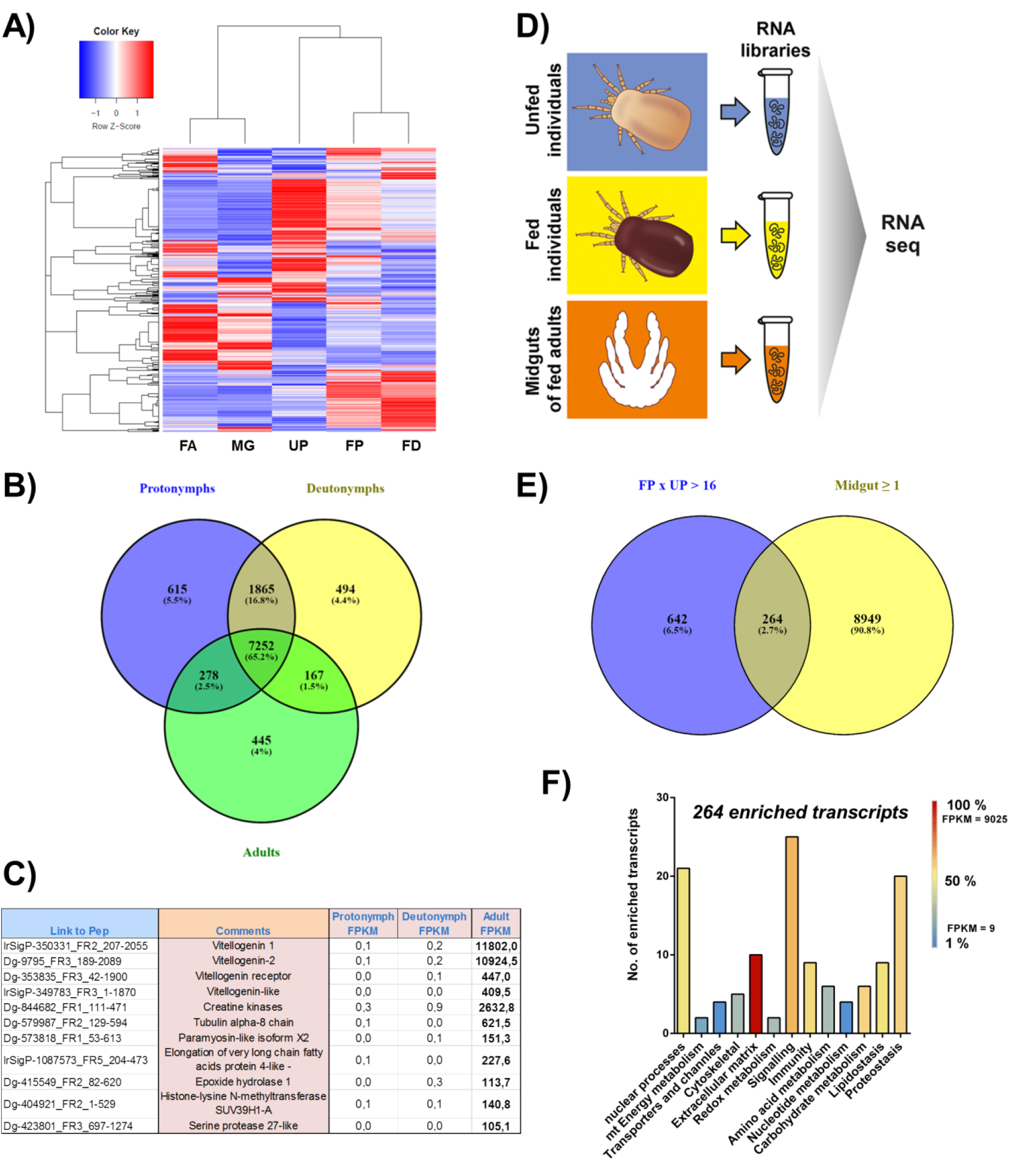
Overview of transcripts encoding developmental stage-specific and digestion-related protein products. **A)** Heat map of RNA-seq transcriptome analysis for 12,484 selected transcripts that had a significant fold change of 16 or higher according to edgeR analysis. UP - unfed protonymph, FP - fed protonymph, FD - fed deutonymph, FA - fed adult, MG - midgut. **B)** Venn diagrams show the stage-specific transcriptomic idiosyncracies as well as the transcriptomic core shared across ontogeny; individual accession IDs are available as Supplementary Data S2. **C)** The table shows top transcripts enriched in transcriptomes of adult mites. These were filtered by >16 × FPKM values in the transcriptome of adults over transcriptomes of both mite juvenile stages, protonymphs and deutonymphs; and E value < e^-80^; protein ID’s are available as Supplementary Data S3. **D)** Depiction of the experimental set up to pin down blood digestion associated encoding transcripts. **E)** Venn diagrams show an overlap of composition with transcripts enriched in fed protonymphs ≥ 16 × over unfed protonymphs and with transcripts of FPKM ≥ 1 in midguts. **F)** The bar graph shows protein classes encoded by individual transcripts enriched by blood-feeding. Transcripts were sorted according to encoded protein class. The number of transcripts from each subclass and their FPKM values are shown. Individual accession IDs are available as Supplementary Data S4.

### Mining the transcriptome for homologous transcripts encoding proteins known to be involved in nutrient sensing, blood digestion, haem and iron biology

#### 1-1) nutrient sensing, digestion, signal transduction, and vitellogenesis

Blood-feeding of arthropods triggers the expression of vitellogenins, precursors of the major egg yolk phosphoglycolipoprotein vitellin ^16^. The interplay between nutrient sensors and initiation of vitellogenesis is well documented in insects ^17^, where the insulin/IGF signaling (IIS) and target of rapamycin (TOR) represent key pathways facilitating the systemic communication between nutrient sensing and reproductive tissues ^18,19^. While the TOR-mediated nutrient-sensing pathway has been partially documented in ticks^20-22^, its description in blood-feeding mites is unknown. Here, we have mined the *D. gallinae* transcriptome and identified (E value < 1e^-50^) individual components of the nutrient sensing and signaling pathway (**Fig. 3A**; Supplementary Data S5). The pathway is initiated by sensing a diversity of building blocks released from digested biomass, both within and outside of the lysosome, as upstream signals ^23^. These activate the master regulator mTORC1 kinase, which gets docked at the lysosomal surface to transduce the nutrient-status signal and stimulate anabolic processes such as vitellogenesis ^22^. To determine whether the *D. gallinae* mites convey feeding-status through a TOR-mediated transduction cascade, thus regulating its anabolism and reproduction, we have used an *ex vivo* membrane feeding system ^24^ to feed adult *D. gallinae* mites with chicken blood supplemented with Torin2, an inhibitor of TOR complexes ^25^. The resulting data show that Torin2 causes clear dose-dependent post-blood-feeding lethality (**Fig. 3B**), thwarting the reproduction of mites.

**Figure 3.**
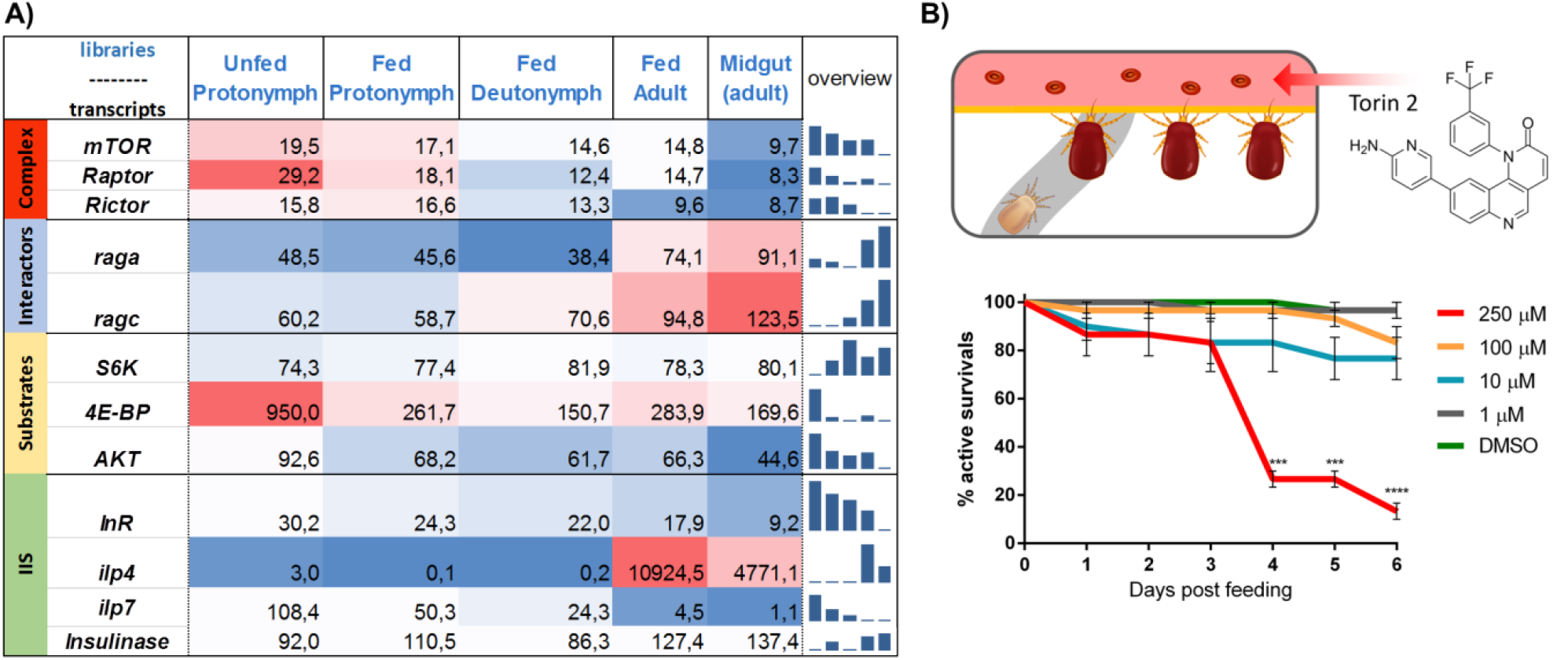
Overview of proteins implicated in nutrient status signaling pathways. **A)** Mapping of nutrient sensing and storage, and TOR-mediated signaling pathway in *D. gallinae* developmental stages, using *C. elegans* homologues as queries ^26^, FPKMs are shown. Accession numbers are available as Supplementary Data S5. **B)** Survival plot of mites exposed to Torin2 inhibitor, through an *ex vivo* membrane feeding system (illustrated on top). Each value is derived from 10 engorged mites in the feeding chamber; Mean and SEM from three independent chambers are shown from two independent experiments. Statistical t-test indicate *** p = 0.0001 and **** p < 0.0001.

#### 1-2) proteolysis

Production of vitellogenins is conditioned by the availability of the basic building blocks - amino acids. In obligate blood-feeders, these ultimately originate from host blood proteins after their catabolic processing by proteolytic enzymes (proteases). Midgut proteases thus represent a major enzymatic suite that enables swift blood processing and a short life cycle turn-around time of *D. gallinae* mites. Transcripts encoding proteases are indeed over-represented in transcriptomes of fed vs. unfed protonymphs and are highly abundant within the midgut-specific transcriptomes (Supplementary Data S4), confirming the importance of these enzymes for the biology of the *D. gallinae* mites. To identify transcripts encoding individual intestinal proteolytic enzymes that catalyse the hydrolysis of host blood proteins, we mined the stage-specific transcriptomes and performed a detailed two-pronged comparative analysis consisting of (*i*) libraries of fed / unfed mite stages, and (*ii*) the midgut / whole body libraries. Cysteine proteases were found to be encoded by the majority of identified reads, accounting for ∼62% of all protease-encoding reads (**Fig 4A**). Transcripts encoding Cathepsin L2 / L3 / L5 account for 80% of all cysteine protease-encoding reads (**Fig. 4B**). These three transcripts displayed a positive ratio of abundance in midgut transcriptomes over the whole body (**Table 1**), and Cathepsin L5 also had a clear ratio of abundance in transcriptomes of fed over unfed mites (**Table 1**), indicating its direct involvement in blood-digestion. The primary sequence analysis revealed that while *Dg*CathL5 shares little homology in the Inhibitor_29 domain with arthropod homologues (18.1 – 33.9 % identity), it displays high homology across the catalytic C1 domain with arthropod homologues (61.5 – 70.1 % identity), including the *Ixodes ricinus* tick Cathepsin L1 (*Ir*CL1) (**Fig. 4C**). *Ir*CL1 is known to display clear haemoglobinolytic and albuminolytic activities, which peak at pH 3.5, indicating its acidic mode of operation within the endolysosomal vesicles of tick gut cells ^27^. It is, therefore, reasonable to speculate that the engagement of the *D. gallinae* Cathepsin L proteases might serve to analogous biological process of host blood protein digestion, possibly also within intracellular acid-lysosomes. However, in contrast to ticks ^28^ and parasitic helminths ^29^, which integrate Cathepsin-B-like enzymes as major components of their multienzyme intestinal proteolytic complexes for blood protein digestion, we have not identified any cathepsin-B homologues in our *D. gallinae* transcriptomes.

**Table 1.** A list of putative digestive proteases and expression values (FPKM) of their mRNA transcripts across studied libraries derived from individual developmental stages of *D. gallinae*.

| Protease | isoenzyme type | Unfed Protonymph | Fed Protonymph | Deutonymph | Adult | Adult midgut | Fed / Unfed | Midgut / Whole Body | RNAseq ID | Protein ID |
| --- | --- | --- | --- | --- | --- | --- | --- | --- | --- | --- |
| Serine Proteases | 1. chymotrypsin | 1,60 | 12,25 | 36,46 | 16,14 | 84,26 | 7,66 | 5,22 | Dg-651195, IrSigP-651191, Dg-651193, Dg-404258, IrSigP-651189 | MBD2881020.1 |
|  | 2. anionic trypsin | 1,24 | 23,91 | 38,00 | 11,60 | 5,34 | 19,28 | 0,46 | Dg-404994 | MBD2877379.1 |
|  | 3. chymotrypsin-like elastase | 5,53 | 21,16 | 12,51 | 18,37 | 44,99 | 3,83 | 2,45 | IrSigP-607270 | MBD2881713.1 |
|  | 4. chymotrypsin-like elastase | 512,79 | 46,02 | 23,34 | 83,31 | 181,47 | 0,09 | 2,18 | IrSigP-76325, IrSigP-355347 | MBD2878320.1 |
|  | 5. stubble-like serine protease, trypsin | 0,06 | 17,89 | 23,88 | 20,77 | 6,74 | 298,17 | 0,32 | IrSigP-387896 | MBD2880678.1 |
|  | 6. stubble-like serine protease, trypsin | 0,12 | 9,57 | 11,95 | 12,57 | 3,86 | 79,75 | 0,31 | Dg-615055 | MBD2879484.1 |
|  | 7. chymotrypsin | 429,29 | 161,08 | 271,43 | 116,19 | 396,48 | 0,38 | 3,41 | IrSigP-417374, Dg-417373 | MBD2877373.1 |
|  | 8. chymotrypsin | 734,01 | 200,66 | 93,83 | 116,07 | 274,78 | 0,27 | 2,37 | IrSigP-622345 | MBD2875824.1 |
|  | 9. chymotrypsin | 569,52 | 639,41 | 682,02 | 503,12 | 1119,46 | 1,12 | 2,23 | IrSigP-997083 | MBD2875973.1 |
|  | 10. chymotrypsin | 151,03 | 46,93 | 39,39 | 24,88 | 60,77 | 0,31 | 2,44 | Dg-602081, Dg-602079 | MBD2879331.1 |
|  | 11. Prolyl oligopeptidase | 64,41 | 188,39 | 170,98 | 434,62 | 282,31 | 2,92 | 0,65 | Dg-1021459, IrSigP-1081927 | MBD2887264.1 |
|  | 12. putative serine protease K12H4.7 | 252,96 | 223,32 | 374,98 | 359,53 | 942,22 | 0,88 | 2,62 | IrSigP-351248, IrSigP-351247 | MBD2884270.1 |
| Cysteine Proteases | 1. cathepsin L1 | 136,58 | 31,43 | 7,07 | 12,84 | 19,49 | 0,23 | 1,52 | Dg-632847, IrSigP-632845 | MBD2879342.1 |
|  | 2. cathepsin L2 | 589,50 | 635,27 | 974,33 | 876,86 | 2250,31 | 1,08 | 2,57 | IrSigP-354374, IrSigP-354370 | MBD2878524.1 |
|  | 3. cathepsin L3 | 114,60 | 263,38 | 488,05 | 478,49 | 1678,34 | 2,30 | 3,51 | IrSigP-413246, IrSigP-10997, IrSigP-413245 | MBD2885740.1 |
|  | 4. cathepsin L4 | 267,69 | 119,88 | 155,36 | 171,78 | 429,18 | 0,45 | 2,50 | IrSigP-349742, IrSigP-349741 | MBD2885566.1 |
|  | 5. cathepsin L5 | 0,61 | 102,01 | 436,16 | 720,11 | 2217,24 | 167,23 | 3,08 | Dg-417098, Dg-417097 | MBD2876666.1 |
|  | 6. cathepsin L6 | 282,77 | 96,61 | 116,45 | 72,91 | 115,77 | 0,34 | 1,59 | IrSigP-1037340 | MBD2884871.1 |
|  | 7. legumain | 98,24 | 10,66 | 12,39 | 13,07 | 45,59 | 0,11 | 3,49 | IrSigP-439085, Dg-439086 | MBD2885742.1 |
|  | 8. legumain | 157,40 | 191,88 | 279,50 | 163,61 | 544,66 | 1,22 | 3,33 | Dg-445098 | MBD2886063.1 |
|  | 9. legumain | 43,35 | 64,32 | 126,99 | 55,15 | 187,81 | 1,48 | 3,41 | IrSigP-535196 | MBD2885743.1 |
|  | 10. legumain | 9,55 | 80,56 | 170,95 | 84,44 | 241,64 | 8,44 | 2,86 | IrSigP-379957 | MBD2885568.1 |
| Aspartyl Proteases | 1. Cathepsin D | 74,33 | 115,50 | 138,36 | 87,39 | 350,90 | 1,55 | 4,02 | Dg-519818, IrSigP-476369, IrSigP-519820, IrSigP-519817 | MBD2876748.1 |
|  | 2. Cathepsin D | 198,87 | 98,69 | 144,50 | 93,47 | 337,65 | 0,50 | 3,61 | Dg-370446, IrSigP-808266 | MBD2879957.1 |
|  | 3. Cathepsin D | 186,75 | 353,83 | 618,02 | 434,81 | 916,66 | 1,89 | 2,11 | IrSigP-16089, Dg-1042377, Dg-1042375 | MBD2876189.1 |
| Metalloproteases | 1. ADAM-like Zinc Metalloprotease | 5,21 | 18,14 | 17,03 | 28,54 | 9,30 | 3,48 | 0,33 | IrSigP-408150, Dg-408153, IrSigP-408144 | MBD2885173.1 |
|  | 2. Zn-dependent nardilysin-like MP | 3,87 | 11,83 | 11,30 | 10,97 | 8,54 | 3,06 | 0,78 | Dg-524874, Dg-524873 | MBD2876667.1 |
|  | 3. peptidase M48C Oma1 (mitochondrial MP) | 11,86 | 34,32 | 22,66 | 14,51 | 13,90 | 2,89 | 0,96 | Dg-514443 | MBD2886064.1 |
|  | 4. M10 heme and peptidoglycan binding MP | 2,23 | 16,85 | 5,42 | 0,00 | 0,00 | 7,56 | 0,00 | Dg-443083, IrSigP-444189, IrSigP-444182, IrSigP-444184 | MBD2887372.1 |
|  | 5. M13 metalloprotease | 15,85 | 9,23 | 5,87 | 5,10 | 17,45 | 0,58 | 3,42 | Dg-566365 | MBD2875907.1 |
|  | 6. Zinc metalloprotease | 18,42 | 23,54 | 33,98 | 11,29 | 37,84 | 1,28 | 3,35 | Dg-586096 | MBD2877380.1 |

**Figure 4.**
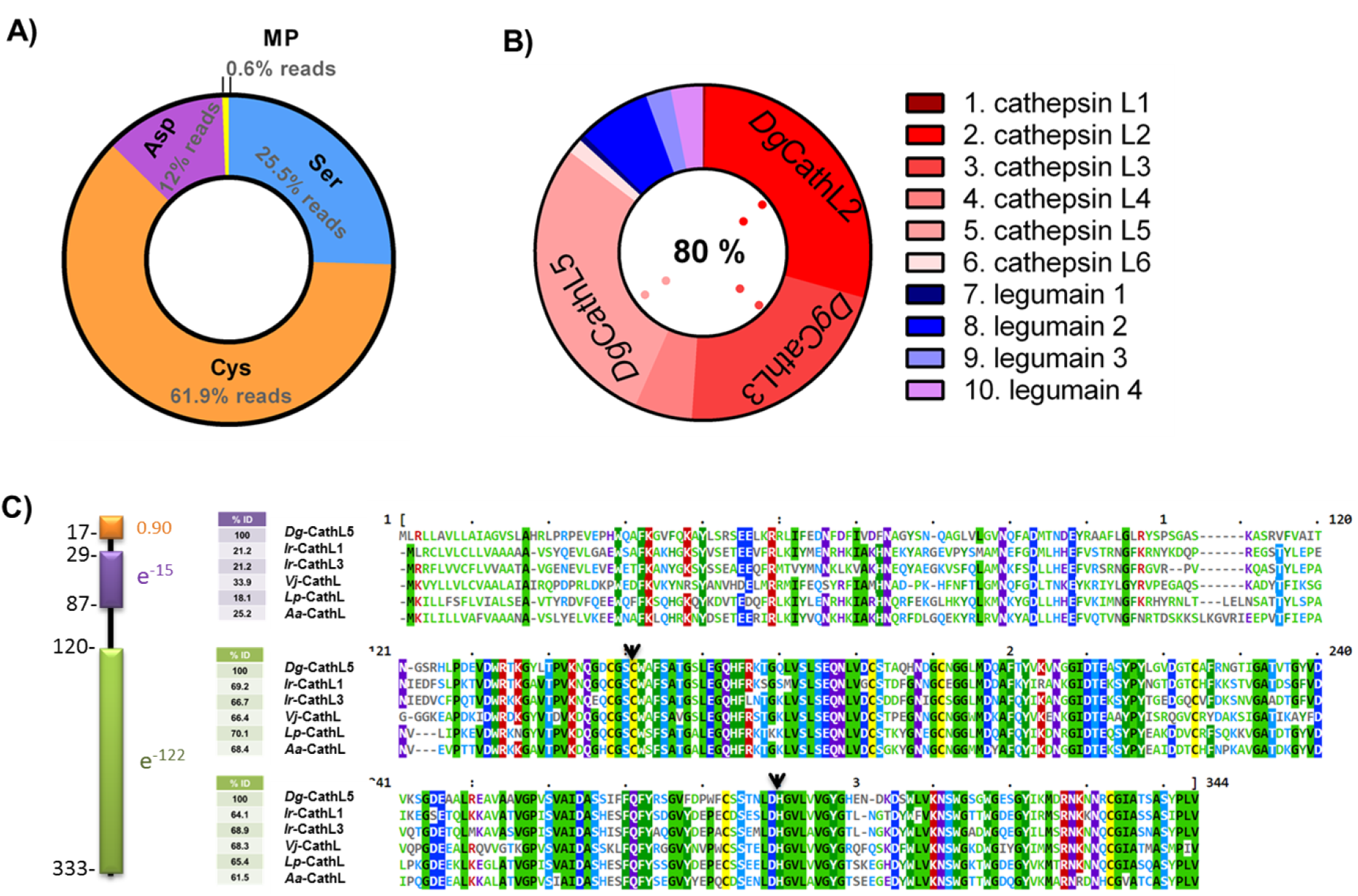
Proteases-encoding transcripts within the *D. gallinae* transcriptomes. **A)** A pie chart of the partition of individual protease families identified in the *D. gallinae* transcriptome; Asp, Cys, Ser - Aspartyl, Cysteine, Serine protease. **B)** A pie chart of individual enzymes identified within the family of encoded cysteine proteases in the *D. gallinae* transcriptome. **C)** Amino acid alignment of arthropod cathepsin L cysteine proteases, homologous to *D. gallinae* cathepsin L5. Schematic view of protein domain arrangements. Signal peptide (orange), predicted by SignalP ^30^, is constituted by the first 17 amino acids (probability = 0.90). CDD domain prediction ^31^ identifies the domain Inhibitor_I29 (violet) with Evalue of e^-15^ and the peptidase C1 domain (green) with Evalue of e^-122^. The amino acid alignment of arthropod cathepsin L homologues includes sequences of: Dg, *Dermanyssus gallinae* (ID: Dg-417098_FR1_205-573); Ir, *Ixodes ricinus* (CathL1 gbID: EF428205.1; CathL3 gbID: MH036745.1); Vj, *Varroa jacobsoni* (ID: XP_022703927.1); Lp, *Limulus polyphemus* (ID: XP_022236235.1); Aa, *Aedes aegypti* (XP_001655999.2). Conserved cysteines are highlighted in yellow. Clustal Omega (EMBL-EBI) was used for multiple aligning of sequences and the alignment was visualised by M-View (EMBL-EBI) ^32^. Active site amino acid residues, cysteine and histidine, are indicated by a black arrow.

For a more comprehensive overview, we have analysed the RNA-seq datasets of *D. gallinae* mites for the presence of mRNA encoding digestive proteolytic enzymes (**Table 1**).

Among the 10 cysteine proteases, we have identified 6 cathepsin-L-like molecules and 4 legumains (asparaginyl endopeptidases). Cathepsin L5 (*Dg*CathL5) is strongly upregulated by nymphal feeding (167 fold) and appears to be primarily expressed in the midgut in adult *D. gallinae*. High read numbers obtained in adult midguts also indicate that the cathepsins L 2, 3, 4 are directly involved in host blood digestion in the mite midgut. In contrast, cathepsin L1, as well as one of the legumains (isoenzyme 7) **(Table 1)**, appear to be feeding-downregulated and their total mRNA loads do not indicate their direct involvement in blood meal digestion. Multiple legumain isoenzymes have been previously demonstrated in the *D. gallinae* related spider mite *Tetranychus urticae* ^33^ *and ticks of the genus Ixodes* have been shown to possess an unusually wide range of functional and non-functional legumain isoforms in their genomes ^34^. All of the four identified *D. gallinae* legumains are primarily expressed in the midgut of adult *D. gallinae*. One of the four identified legumain mRNA, the isoenzyme 9 in **Table 1**, encodes a pseudoprotease that has histidine and cysteine active site residues replaced by leucine and serine, respectively. The isoenzyme 10 in **Table 1** is upregulated by nymphal feeding (8.5 fold) which indicates its possible association with blood digestion.

Of interest are also three aspartyl proteases of the cathepsin-D type (pepsin family) that clearly display midgut-specific expression (**Table 1)**. Their high levels of transcripts indicate the direct involvement of these proteases in host protein digestion, analogously to other Acari such as *Sarcoptes scabei* ^35^ and ticks ^36,37^. Of interest from serine proteases are two stubble-like serine proteases (isoenzymes 5 and 6 in **Table 1**), whose expression is strongly upregulated by blood-feeding (by 288 and 78 fold, respectively). Their total quantity of mRNA reads remains relatively low, which may indicate their potential role in molecular signaling rather than direct involvement in catabolism of host blood proteins. None of the several metalloprotease-(MP)encoding mRNAs with responsive expression were present in large amounts, indicating their role in regulation and signaling pathways rather than in protein metabolism. Of interest are M13 and zinc metallopeptidases (MP types 5 and 6 in **Table 1**), which are predominantly expressed in the midgut (**Table 1)**. Additionally, all identified protease isotypes were further analysed for the presence of a signal peptide. The full primary protein sequences, including the identical and deduced allelic protease variants with an indicated signal peptide, active site residues, and where applicable, also corrected or deduced N and C termini, are listed in Supplementary Data S6.

Blood feeding has evolved several times in Acari ^38^. The haemoglobinolytic apparatus was suggested to be shared among phylogenetically related blood-feeding parasites and highlighted as promising anti-parasite targets ^28^. Although ticks and dermanyssoid mites belong to the same order Acarina (class Arachnida), and both represent blood-feeding ectoparasites, the data obtained here support the hypothesis of their independent appearance of blood-feeding. This is substantiated by their clear differences in digestive proteolytic machinery. While both hard (Ixodes ricinus) and soft ticks (*Ornithodoros moubata*) rely on cathepsin B as the major hemoglobinolytic peptidase ^39^, we have not identified any cathepsin B-like protease encoding transcripts. Instead, the intestinal proteolytic machinery for host blood digestion seems to rely on cathepsin L -like cysteine proteases as the major components, although the roles of legumains, cathepsin D, and other components might complement the rest of the tick intestinal proteolytic machinery. This finding confirms the earlier observation, showing a clear capacity of *D. gallinae* homogenate to digest haemoglobin at an acidic pH, with the activity being inhibitable by E-64 and pepstatin A, indicating the participation of cysteine and aspartic proteases ^40^.

In summary, the less-than-an-hour long on-host feeding ^41^, during which time, mites increase their body weight by approx. ten fold ^42^, is followed by rapid off-host digestion. About half of the imbibed erythrocyte content is digested within 4 – 8 hours in the close relative dermanyssoid mite *Ornithonyssus sylviarum* ^43^. *Similarly to ticks and acariformes mites* ^*44*^, *the parasitiforme blood-feeding mites (D. gallinae*) ^40^ appear to digest host blood intracellularly using a multi-enzyme complex of mainly acidic endolysosomal proteolytic enzymes, yet lacking any cathepsin B homologue.

#### 1-3) blood digestion - metallostasis

Apart from being protein-rich, host blood is also a rich source of haem- and non-haem iron. Unlike ticks that cannot synthesise haem *de novo* and must take haem from host blood haemoglobin ^45^, *D. gallinae* mites clearly display the capacity to synthesise haem *de novo*. First, we managed to identify stable intermediate products of haem biosynthesis in *D. gallinae* homogenates (**Fig. 5A**) and, second, we identified transcripts encoding a full enzymatic set for haem biosynthesis (**Fig. 5B**). In addition to molecular detection of functional haem biosynthesis (**Fig. 5C**), the fact that unlike ticks (*Ixodes ricinus*) ^45^, *D. gallinae* mites produce colorless eggs (**Fig. 5D**), indicating the absence of haem deposits, provides another layer of evidence for a different biology of haem between ticks and *D. gallinae* mites, despite both being obligatory blood-feeders.

**Figure 5.**
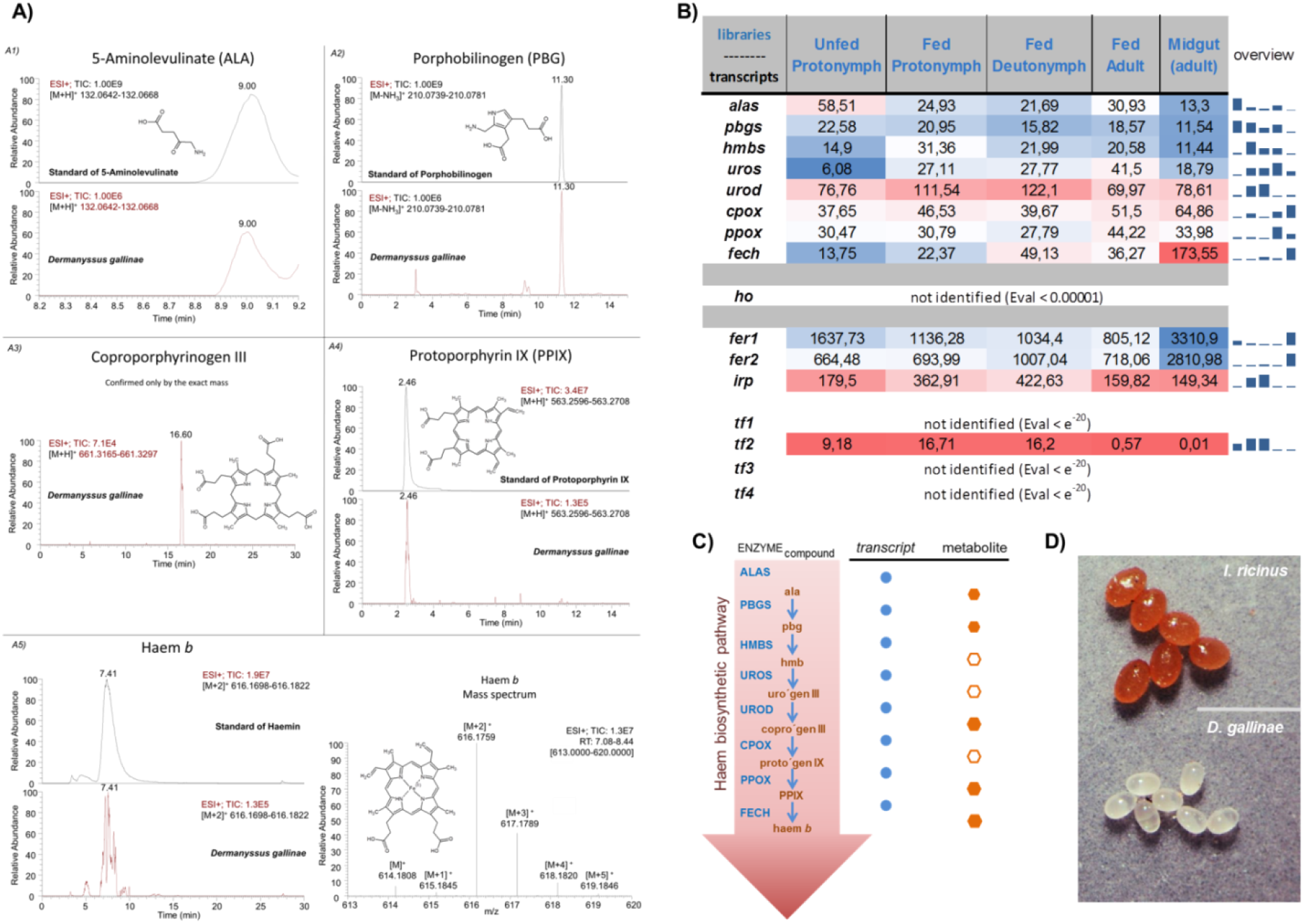
Haem and iron homeostasis-related protein-encoding transcripts within the *D. gallinae* transcriptomes. **A)** The LC/MS analysis of Haem *b* and its precursors in the naive non-fed stages of *Dermanyssus gallinae*. **A1)** Reconstructed chromatogram for 5-Aminolevulinate [M+H]^+^ 132.0642-132.0668, first for the standard and second for the *D. gallinae* sample. **A2)** Reconstructed chromatogram for Porphobilinogen [M-NH3]^+^ 210.0739-210.0781, first for the standard and second for the *D. gallinae* sample. **A3)** Reconstructed chromatogram for Coproporphyrinogen III [M+H]^+^ 661.3165-661.3297 for *D. gallinae* sample. Confirmed only by the exact mass. **A4)** Reconstructed chromatogram for Protoporphyrin IX [M+H]^+^ 563.2596-563.2708, first for the standard and second for the *D. gallinae* sample. **A5)** Reconstructed chromatogram for Haem *b* [M+2]^+^ 616.1698-616.1822, first for the standard (Haemin) and second for the *D. gallinae* sample and mass spectrum of Haemin for explanation of the diagnostic mass [M+2]^+^. **B)** Table of proteins participating in haem or iron homeostasis, together with their expression values (FPKM) of their mRNA transcripts across libraries derived from individual developmental stages of *D. gallinae*. Accession numbers are available as Supplementary Data S8. **C)** A schematic of haem biosynthesis, which identifies trancripts and metabolites of the pathway. **D)** A photographic image of eggs of *Ixodes ricinus* ticks and *D. gallinae* mites, note the colour difference, indicating the presence/absence of deposits of maternal haem.

Similarly to ticks and other mites ^45^, *D. gallinae* mites do not code for haem oxygenase, a haem-cleaving enzyme that liberates bioavailable iron (**Fig. 5A**). *D. gallinae* mites thus likely acquire iron from non-haem sources, possibly from serum transferrin as ticks do ^45^. Acquired iron is likely stored intracellularly in ferritin 1 (Fer 1). We have identified a single homologue of D. *gallinae fer1* transcript with a conserved iron-responsive element in the 5’UTR region ^46^ (Supplementary Data S7) and clear midgut-enriched expression (**Fig. 5A**), both supporting the participation of Fer1 in storage of dietary iron in the mite midgut. While insects distribute acquired iron somatically, mainly by transferrin 1 ^47^, ticks distribute dietary iron somatically through a secretory type of invertebrate ferritin, Fer 2 ^48^. We have identified in the *D. gallinae* transcriptome a single homologue of tick secretory Fer2, *dg*-*fer2*, with clear midgut-enriched expression, but failed to identify a homologue of insect Transferrin 1 (**Fig. 5A**, Supplementary Data S8), indicating Ferritin 2-based somatic distribution of iron in the *D. gallinae* mites.

Vitellogenins, precursors of vitellins, are the predominant lipoproteins expressed in fat bodies and midguts, and then transported into developing ovaries ^45^. While tick vitellins also bind haem, acquired from host blood haemoglobin, vitellins from other organisms do not ^49^. In ticks, unlike other haemolymph lipoproteins ^50^, true vitellogenins are solely expressed in mated engorged females ^45^. Using the same criterion, we have identified, within the *D. gallinae* transcriptome of adult mites, two vitellogenin transcripts encoding homologues of tick vitellogenin 1 and vitellogenin 2 ^45^, and also an additional vitellogenin transcript, which seems to be unique for Dermanyssoidea mites, denoted as Vg1-like (**Fig. 6**, Supplementary Data S9). All three transcripts clearly displayed enhanced levels within transcriptomes of adult mites and with virtually no mRNA present in the transcriptomes of juvenile stages (**Fig. 2C**). Phylogenetic analysis of arthropod vitellogenins produced several vitellogenin clades reflecting their arthropod taxonomic grouping, i.e., parasitiform ticks and mites, acariform mites, spiders, scorpions, horseshoe crabs, insects, and crustaceans (**Fig. 6**). The acarids split into two well-supported clades containing homologues from the Parasitiformes (ML/BI = 95/1.00) and the Acariformes (ML/BI = 84/0.98). Parasitiform sequences clustered within the well-supported Vg1 and Vg2 lineages (both ML/BI = 100/1.00), each comprising two tick and two to three mite homologues. The *D. gallinae* mite Vg1 (transcript IrSigP-350331_FR2_207-2055, Protein ID: MBD2876257.1), Vg1-like (transcript IrSigP-349783_FR3_1-1870, Protein ID: MBD2876000.1) and Vg2 (Dg-9795_FR3_189-2089, Protein ID: MBD2876256.1) grouped with their respective mite vitellogenin orthologues within the corresponding Vg lineages (**Fig. 6**).

**Figure 6.**
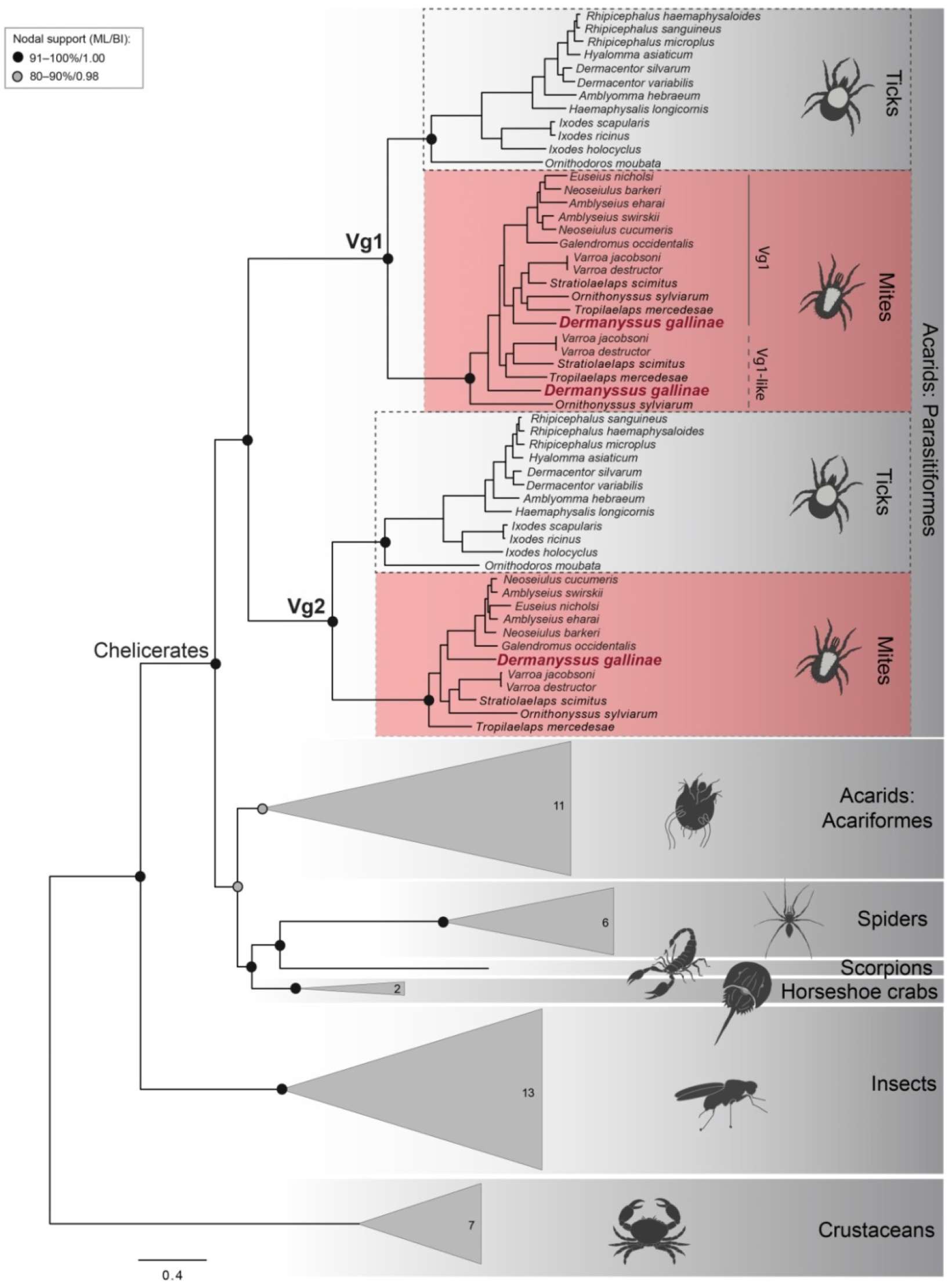
Phylogenetic relationships of arthropod vitellogenins (Vg) with a focus on their evolution in mites and ticks. Maximum likelihood phylogeny of 94 arthropod vitellogenin amino acid sequences showing the positioning of three *Dermanyssus gallinae* vitellogenin homologues within the Vg1 and Vg2 lineages. Crustacean vitellogenins were used as an outgroup. Nodal supports at the main nodes are represented by maximum likelihood bootstrap values and Bayesian inference posterior probabilities, respectively. For simplification, the homologues from non-parasitiform taxa were collapsed into triangles; numbers inside the triangle indicate the number of sequences included in each clade. For GenBank accession numbers, see Supplementary Data S9.

### Catalogue of acaricide targets: Ligand-gated Ion Channels

The majority of commercially available acaricides, as well as insecticides, target invertebrate nervous systems, usually by interaction with an allosteric site of ionotropic receptors, also called ligand-gated ion channels (LGICs) ^51^. Members of this family have a high degree of primary sequence similarity, forming a ligand-binding site for neurotransmitter molecules and an ion-conducting pore. LGICs can be categorised into four major families ^52^, one of which entails a series of validated acaricidal/insecticidal targets, i.e. the pentameric cys-loop family, which shares a highly characteristic motif, a 15-aa cys loop in the protein N-terminus ^53,54^. The cys-loop receptors share a three dimensional modular structure, forming a pentameric pore, which is lined with second membrane-spanning domains of each subunit. The cys-loop receptors are both cation-permeable, such as nicotinic acetylcholine receptors (nAChRs) ^55^ and serotonin receptors, and also receptors that conduct anions, such as γ-amino butyric acid (GABA)-gated ion channels ^56^, glycine receptors, as well as chloride channels gated by glutamate (GluCls), histamine ^57,58^ or zinc ^59^. The last two, histamine- and zinc-gated chloride channels, were not identified in the *D. gallinae* transcriptome. Unlike some pentameric ligand-gated ion channels, homologues of RDL(resistance to dieldrin)-GABA receptors ^51^ and GluCls are unique to invertebrates ^60^. RDL-GABA and GluCl channels can be targeted by negative modulators such as fipronil, or positive modulators, such as ivermectin ^61^. We have mined our transcriptome and reconstructed a phylogenetic tree of cys-loop receptors, having identified (E value < e^-10^, using *Pediculus* sp. ^62^ and *Tetranychus* sp. ^63^, as queries) 29 transcript-encoding members of the cys-loop family (**Fig. 7A**, Supplementary Data S10). The *D. gallinae* transcripts clustered with their respective homologues from arthropods and vertebrates into nine clades and in an additional clade that exclusively united the Acari-specific glutamate gated chloride channel-like proteins. We have identified four transcripts encoding glutamate-gated chloride channels, two of which display midgut-specific expression (**Fig. 7B**). Despite the fact that the Dg-565248_FR2_220-775 transcript encodes a truncated N-terminus (Protein ID: MBD2887064.1), thus lacking a predicted signal sequence, all transcripts shared a core of GluCl channels with the Cys-Loop region and mostly four transmembrane domains (Supplementary Data S11). The amino acid sequence analysis of GluCl channels demonstrates conservation of most of the amino acid residues required for ivermectin association with the *C. elegans* GluCl alpha channel ^64^ (Supplementary Data S11). The assembly of the four GluCl homologues is supported by multiple reads, except for the encoded C-terminus of the IrSigP-571488 transcript, which is composed of uniquely assigned reads (Supplementary Data S12). To confirm sensitivity of *D. gallinae* mites to acaricides targeting GluCl channel(s), as exemplified by Fipronil, Ivermectin, and Fluralaner, we experimentally determined the dose-dependent survival after microinjection of the respective compounds into adult mites (**Fig. 7C**). While Ivermectin and Fluralaner elicited clear lethal effects within a couple of days upon microinjection of a 10µM solution, Fipronil caused only negligible lethality even when injected at 100 µM (**Fig. 7C**). Looking at the midgut/whole body ratio, we filtered out four more transcripts with a midgut-enriched presence of mRNA transcripts (**Fig. 7D**), one encoding the pH-sensitive chloride channel protein and three encoding Acari-specific chloride channel-like proteins. While we have no knowledge on the Acari-specific chloride channels, an increasing body of work is available on pH-sensitive chloride channels in other invertebrates. Interestingly, the pH-sensitive chloride channel from *Bombyx mori* is sensitive to Ivermectin, but non-responsive to Fipronil ^65^, which is reminiscent of our data from viability assays upon the microinjection of agonists. This may suggest that pH-dependent chloride channels could be key targets for channel agonists like Ivermectin in eliciting acaricidal activity.

**Figure 7.**
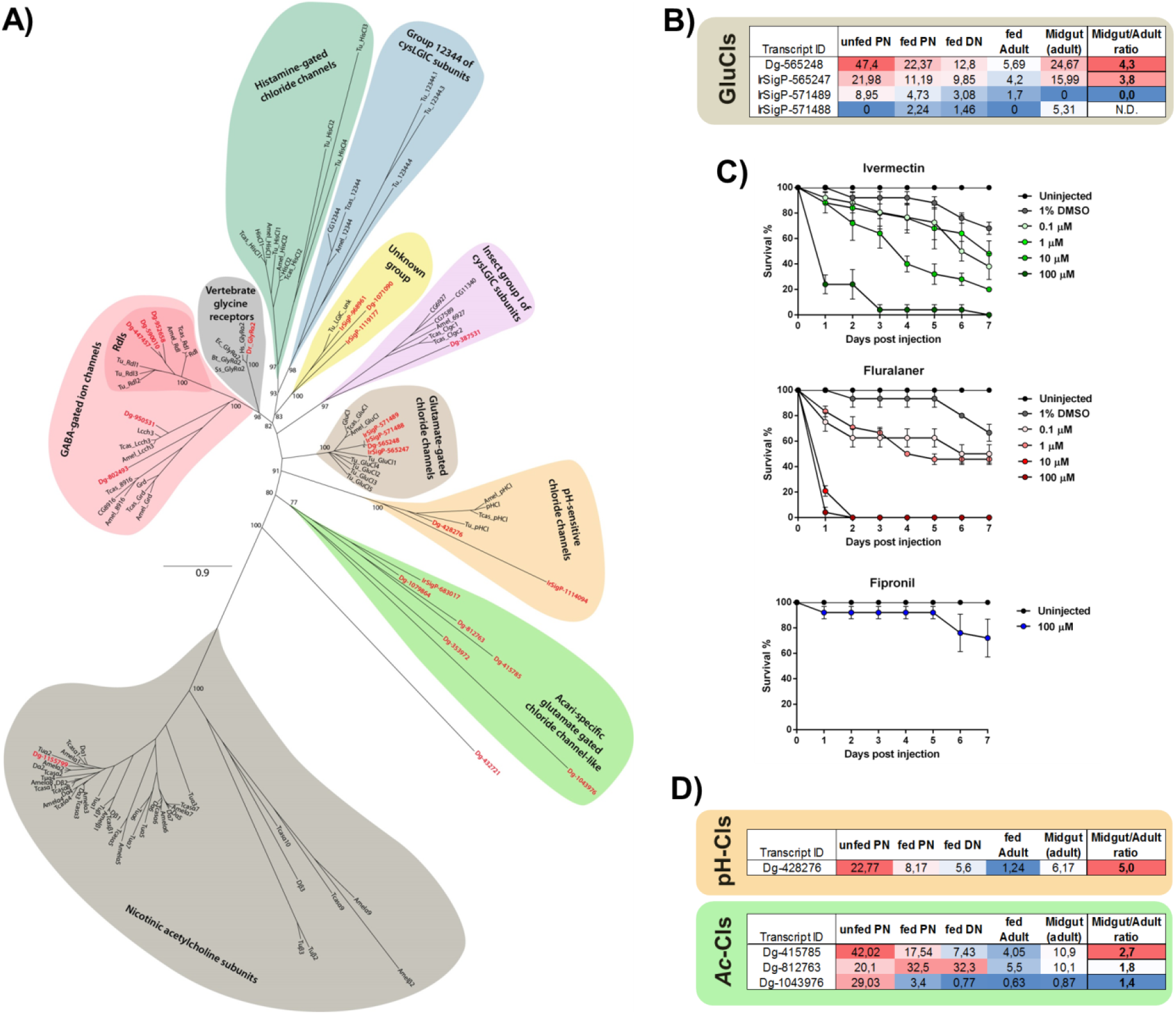
Glutamate gated chloride channels as acaricidal targets. **(A) Maximum likelihood phylogenetic tree of the arthropod and vertebrate cysLGIC subunit protein sequences**. The bootstrap supports above 50% are shown at the main nodes. *Dermanyssus gallinae*: Dg and IrSig (indicated in red), *Tetranychus urticae*: Tu, *Drosophila melanogaster* (D or other), *Apis mellifera* (Amel), *Tribolium castaneum* (Tcas), *Homo sapiens* (Hs), *Danio rerio* (Dr), *Bos taurus* (Bt), *Equus caballus* (Ec), *Sus scrofa* (Ss), and *Gallus gallus* (Gg). For accession numbers see Supplementary Data S10. **(B)** FPKM values of individual GluCls and their midgut/whole body ratio of adult mites. **(C)** Dose-dependent viability assays after microinjection of GluCls agonists dissolved in DMSO and diluted to 1% of the solvent. Each data point represents a mean, and SEM of n = 25 mites per tested concentration. **(D)** FPKM values of individual Acari-specific (Ac) and pH-dependent (pH) chloride channels (Cls) and their midgut/whole body ratio of adult mites.

### Mining of pathogen recognition repertoire, immune signal transduction pathways, RNAi, and immune elicitors

Innate immunity is highly conserved in metazoans spanning both vertebrates and invertebrates ^66^. The humoral part of the invertebrate immunity comprises the sensing of microbial non-self, which is followed by signal transduction, mainly through Toll and Imd signaling pathways, which operate to some extent independently ^67^. Here, we have mined the *D. gallinae* transcriptome and managed to reconstruct a complete Toll pathway, suggesting full functionality in *D. gallinae* (**Fig. 8**, Supplementary Data S13). The only missing components were Gram-negative bacteria-binding proteins (GNBPs) and the transcription factor DIF. Peptidoglycan recognition proteins (PGRPs) were present in two different variants with uncertain classification. Nine Spätzle proteins and eight Toll receptors were identified. The Imd pathway was significantly reduced, as most components were missing, including IMD and Relish. The retained members identified in the *D. gallinae* mites, which are integrated in the IMD pathway of insects, were Caspase (DREDD, Cysteine-dependent ASPartyl-specific proteASE) and Serine/Threonine Kinase (TAK1 and IKKβ). The absence (not identified in our transcriptome, nor in the genome with an Eval cut off = 0.0001) of a critical component of the pathway, transcription factor Relish, indicates non-functionality of the IMD pathway in the *D. gallinae* mites, similarly to the phylogenetically related mites, Varroa (Parasitiformes; Eval cut off < 10e^-3^) and Metaseiulus (Acariformes) ^68^. Blood-feeding arthropods are notorious vectors of human and animal viruses, some of which cause debilitating diseases. Potent immune responses need to be deployed in blood-feeders in order to sense and regulate their endogenous viral load. Arthropods ^69^, similarly to higher organisms ^70^, are armed with an RNA-based mechanism of anti-viral immunity called RNAi. This pathway is functional against both RNA and DNA viruses in insects ^71,72^. Both *in silico* predictions and experimental validation supports the presence of a functional RNAi pathway in *D. gallinae* ^73^. We have confirmed (**Fig. 8**) the *in silico* analysis ^73^, and managed to identify most of the transcripts encoding RNAi proteinaceous components, with the exception of an R2D2 homologue (Eval cut off < 0.00001; query *Drosophila* R2D2: NP_001285720.1). R2D2, a dsRNA binding protein, is reported to be essential for the loading of siRNAs into effector Ago-RISC complexes ^74^. This is in line with a comprehensive phylogenetic analysis that determines the existence of R2D2 homologues only in orders of winged insects (Pterygota) ^75^. Reconstruction of the ancestral state suggests that R2D2 is a derived feature (autapomorphy) of Pterygota. The absence of R2D2 in primarily wingless insects, as well as in non-insect arthropods (ticks, mites, crustaceans), however, does not necessarily mean that these species lack a functional RNAi pathway. The authors further speculate that it is possible that the corresponding gene from the miRNA pathway (Loquacious) could compensate for R2D2 in these species ^75^.

**Figure 8.**
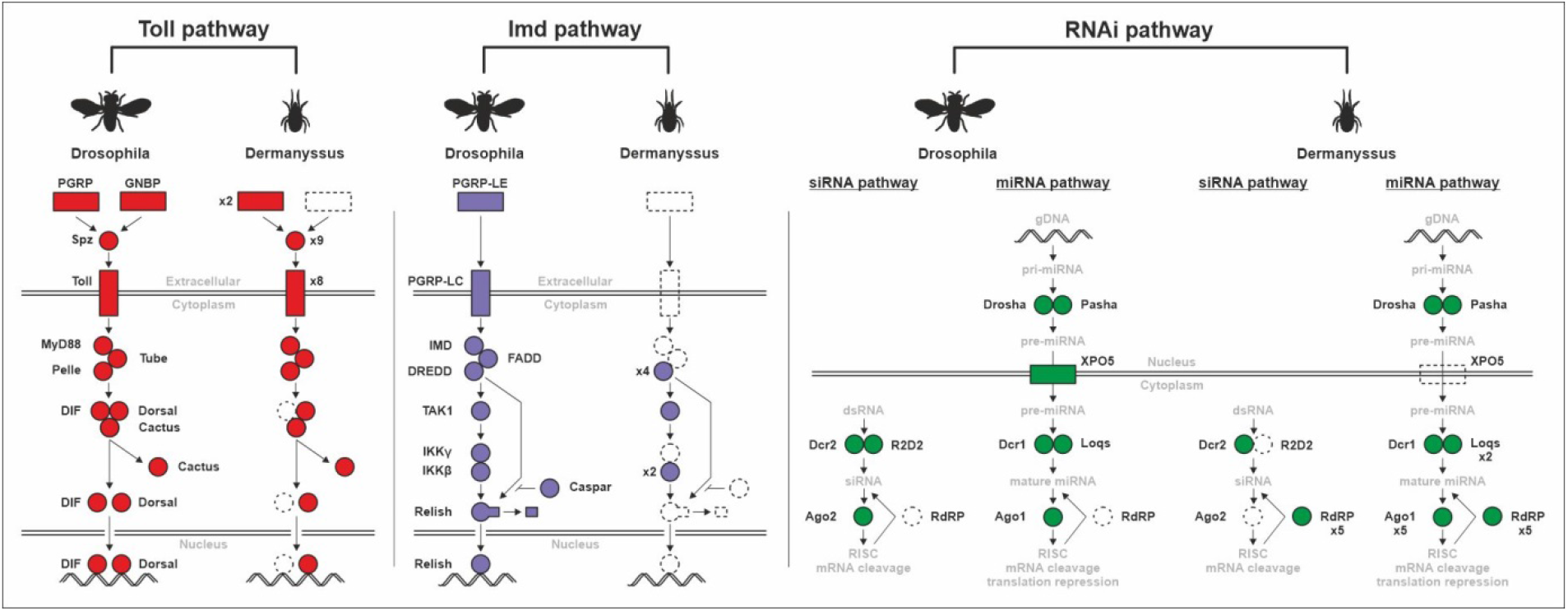
Mining of Toll, IMD, and RNAi pathway in the *D. gallinae* transcriptome. Protein sequences of *Drosophila* or *Tribolium* (Tube) Toll and Imd pathways were used to search in our *D. gallinae* translated protein database by using program Bioedit (Local BLAST, E value 0.1, Matrix BLOSUM62). Conservation of the domains was checked by CD-search (NCBI). Only hits with similar domain structures and an E-value < 10e^-3^ were considered as putative homologues. Accession numbers are available as Supplementary Data S13.

#### i) thioester-containing proteins (TEPs) as the components of the ancestral system of complement

The principal role in cellular and humoral innate immunity of vertebrate as well as invertebrate metazoan organisms is played by the complex complement system that allows for recognition of pathogens, their specific tagging by opsonization, and their elimination via phagocytosis or cell lysis. More than thirty components of the complement system have been identified in higher vertebrates ^76^, but the presence of an ancestral complement system has been evidenced in organisms such as horseshoe crabs that were living on the Earth more than 500 mil. years ago ^77,78^. The central effector molecules of vertebrate and invertebrate complement systems are proteins belonging to the thioester-containing protein (TEP) family, formerly referred to as proteins of the α2-macroglobulin superfamily ^76,79,80^. The invertebrate TEPs are divided into four major phylogenetically distinct groups comprising pan-protease inhibitors of the α2-macroglobulin type (α2M), C3-like complement components (C3), insect-type TEPs (iTEPs), and macroglobulin-complement related (MCRs) ^77,81,82^. Recently available horseshoe crab genomes ^83,84^, together with transcriptome data from a variety of arthropods, reveal that all these major groups of TEPs are present in chelicerates, but C3-like molecules are absent in crustaceans and hexapods and α2Ms were lost in the evolution of some insect lineages such as fruit flies or mosquitoes ^77^. Here, we examined the *D. gallinae* transcriptome with full sequences of well-annotated nine members of the *I. ricinus* TEP family comprising all four groups of invertebrate TEPs ^81,85^. BlastP mining and the following phylogenetic analyses with other selected representatives of invertebrate TEPs revealed that *D. gallinae* possesses one α2-macroglobulin molecule present in at least five splicing variants within the protease-sensitive ‘bait-region’ (DgA2M(sv1-5)(Supplemental Data S14). The diversification of ‘bait-regions’ by alternative splicing extends the portfolio of target proteases inhibited by the mechanism of molecular entrapment ^86^, previously reported for A2Ms from ticks ^87,88^. *D. gallinae* possesses one molecule clearly belonging to the insect-type TEP (*Dg*TEP), one C3-complement like molecule (*Dg*C3) and three molecules belonging to the group of macroglobulin-complement-related proteins (*Dg*MCR-1,2,3) (**Fig. 9**). DgMCR-3 appears phylogenetically quite distant from the other two, *Dg*MCR-1, and *Dg*MCR-2, yet its membership within the MCRs is supported by its stable phylogenetic positioning inside the MCR clade and by the presence of the signature low density lipoprotein receptor domain in the central part of the molecule ^81,85^. Taken together, *D. gallinae* possesses representatives of all major groups of invertebrate TEPs. The FKPM values of respective TEPs showed substantial differences in their expression (Supplemental Data S14).

**Figure 9:**
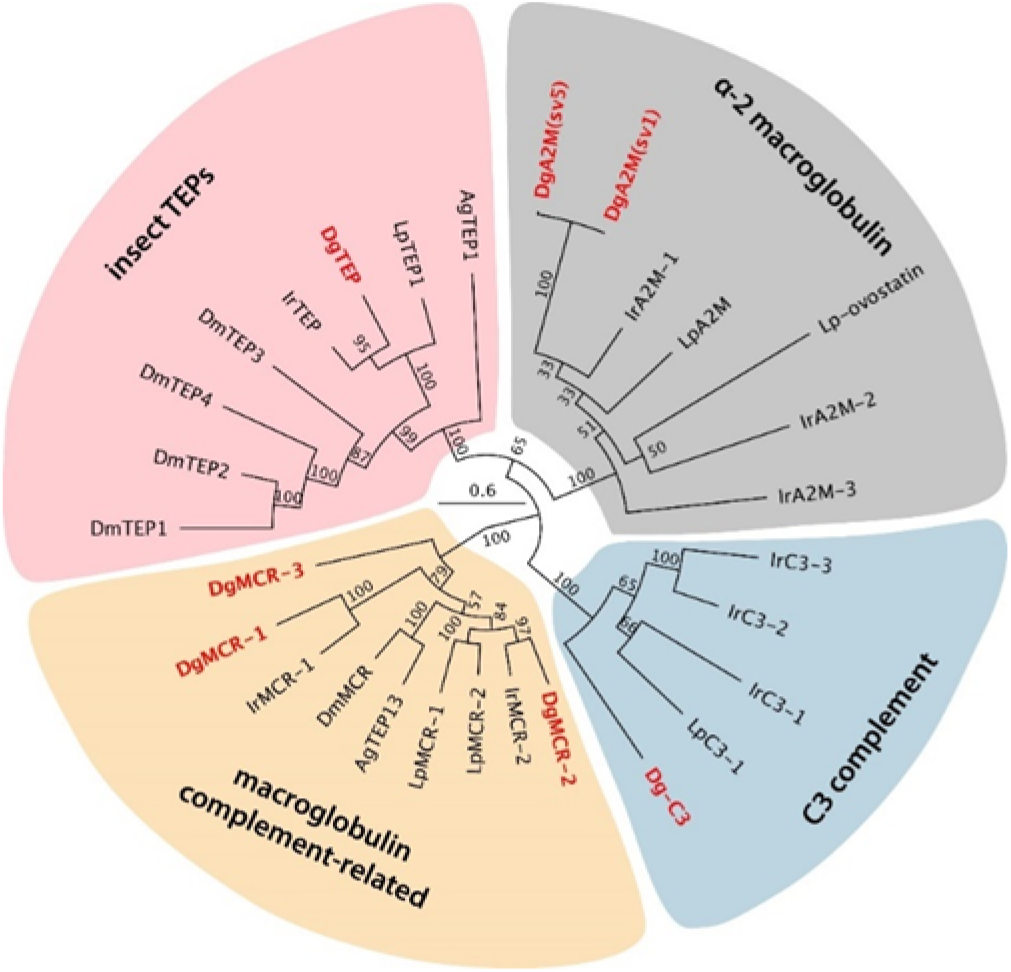
Phylogenetic tree of identified *D. gallinae* members of the thioester-containing proteins (TEP) family with selected well-annotated TEPs from other invertebrates. The tree was reconstructed using the maximum likelihood method based on the alignment of full amino acid sequences (∼1500 residues). Ag – malaria mosquito *Anopheles* gambiae; Dg (in red) – red poultry mite *Dermanyssus gallinae;* Dm – the fruit fly *Drosophila melanogaster*; Ir – the hard tick *Ixodes ricinus;* Lp – the horseshoe crab *Limulus polyphemus*. Numbers at the branches represent bootstrap support values calculated from 1,000 replicates. For respective accession numbers, see the Supplementary Data S14.

#### ii) defensins as the representatives of the effector antimicrobial peptides

Defensins are probably the most widely spread antimicrobial peptides of 3 – 5 kDa, broadly distributed in the animal and plant kingdoms ^89^. We have searched for defensin-like molecules in the *D. gallinae* transcriptome using the data from the tick *Ixodes scapularis* genome that encodes multiple defensin-related molecules, divided into two major families: (i) scapularisins, structurally related to the ancient invertebrate-type defensin and (ii) scasins, which are only distantly related to the canonical defensins ^90^. While we failed to find any transcript related to the *I. scapularis* scasin (Gen Bank EEC18782), we identified several defensin-like molecules related to the scapularisins with the highest homology displayed to scapularisin-6 (GenBank EEC08935) annotated as defensin-2 in the *I. scapularis* genome (VectorBase ISCW005928). Multiple sequence alignments of eight *D. gallinae* transcripts encoding putative defensins with scapularisin-6 (**Fig. 10**) revealed that *D. gallinae* defensins can be divided into two major types: (i) Type I defensins (in blue) lack the canonical furin cleavage motif (RVRR) ^91^ and (ii) Type II defensins (in red) having the furin site conserved. Based on the FPKM values, Type I defensins seem to be much more expressed in all stages (Supplemental Data S15).

**Figure 10:**
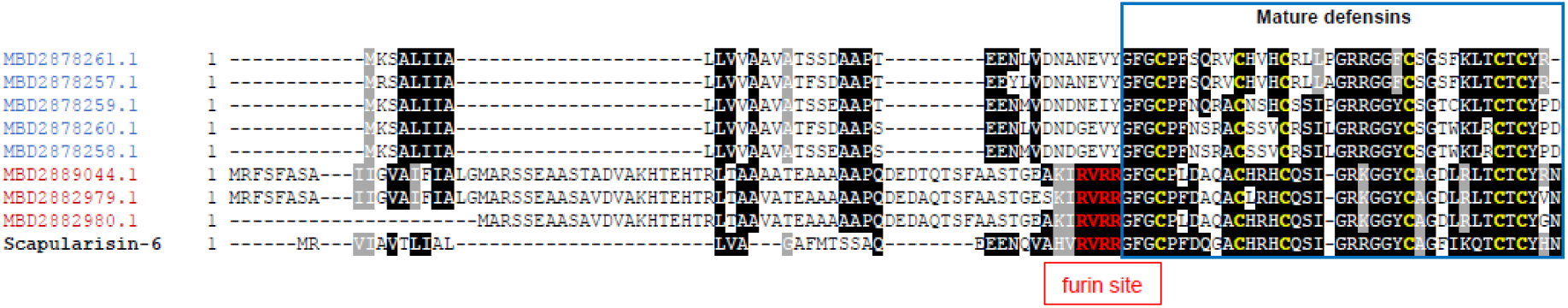
Multiple sequence alignment of identified *D. gallinae* pre-prodefensins with the tick *Ixodes scapularis* scapularisin-6 (GenBank EEC08935). For accession numbers, homology values and FKPM values in *D. gallinae* developmental stages, see Supplemental Table Supplemental Data S15.

### Description of RNA-virome in the *D. gallinae* mites

*D. gallinae* has been linked with several poultry infections, serving as a likely vector of detrimental diseases of hens. In addition, the detection of human viruses such as St. Louis encephalitis virus ^92^, in concert with *D. gallinae* wide host range comprising wild birds and several mammals, including humans, suggest that these mites could be of wider concern for veterinary and medical science as a potential vector for pathogenic agents ^10^. Interestingly, to our knowledge, there are no reports of *D. gallinae* mite-specific viruses. As a part of our aim to characterize the complete RNA component of *D. gallinae*, we oriented our search, using a standard pipeline of arthropod virus discovery, to detect potential viral like sequences in our red mite RNAseq samples. Using as the query, NCBI-refseq virus proteins, we subjected our transcriptome consensus assembly to tblastN searches (e-value < 1e^-5^), resulting in several putative transcripts with significant hits to viral proteins (**Table 2**).

**Table 2.** Expression values (FPKM) of Quaranjavirus genome segments across libraries derived from individual developmental stages of red mites.

| libraries |  |  |  |  |  |
| --- | --- | --- | --- | --- | --- |
| ----- |  |  |  |  |  |
| Quaranjavirus segments | Unfed Protonymph | Fed Protonymph | Fed Deutonymph | Fed Adult | Midgut (adult) |
| *RMQV1-HA* | 101 | 245 | 760 | 110 | 418 |
| *RMQV1-NP* | 261 | 285 | 624 | 135 | 510 |
| *RMQV1-PA* | 21 | 94 | 230 | 43 | 76 |
| *RMQV1-PB1* | 43 | 96 | 245 | 52 | 150 |
| *RMQV1-PB2* | 61 | 93 | 194 | 12 | 116 |

Our attention was caught by a specific hit in our database, that of a 1.5kb transcript with resemblance to the hemagglutinin protein (HA) of Wellfleet Bay virus (WFBV, 33.85% identity, e-value 9e^-96^). WFBV is the causative agent of cyclic large-scale massive bird die-offs ^93^. WFBV is a member of the *Quaranjavirus* genus of negative-sense single-stranded enveloped multisegmented RNA viruses of the family *Orthomyxoviridae*, which includes the influenza viruses. In additional tblastN searches using, as a query, all available quaranjavirus proteins, we were able to detect four further transcripts resembling all the typical core genome segments of orthomyxoviruses. This included a best hit to a 1.8kb transcript of the nucleoprotein (NP) of *Quaranfil quaranjavirus* (QRFV, identity 30.20%, e-value 6e^-57^). QRFV is the type species of the genus *Quaranjavirus*, a virus hosted by ticks, birds and also able to infect humans, as reflected by a fundamental serological study during the 1960s in Egypt, which found that 8% of the local population had neutralizing antibodies to QRFV ^94^. Besides these structural proteins, we found three transcripts ranging from 2.4kb to 2.5kb with best hits to the multipartite RNA dependent RNA polymerase sub unit proteins PB1 (Tjuloc virus, identity 50%, e-value 0), PB2 (QRFV, identity 28.90%, e-value 9e^-94^) and PA (Tjuloc virus, identity 50%, e-value 0). Tjuloc virus was originally isolated from *Argas vulgaris* ticks linked to multispecies birds in Kyrgyzstan ^95^. These transcripts were polished and curated using the filtered reads of each library to generate consensus sequences of each genome segment, supported by a mean coverage ranging from 13.5X to 47.5X. The putative genome segments of expected size were further annotated and, as anticipated, each presented a single ORF in their complementary RNA+ strand coding for the structural and functional proteins of a typical quaranjavirus, flanked by un-translated regions (**Fig. 11** left panel, Supplementary Data S16). The translated polymerase subunit proteins PB1, PB2 and PA presented with conserved domains Flu_PB1 (pfam00602, e-value 3.62e^-60^), Flu_PB2 (5D98_F, e-value 8.2e^-121^), and Flu_PA (4WRT_A, e-value 4.9e^-97^), respectively. The NP showed a Flu_NP domain (3TJ0_B, e-value 7.5e^-81^) and the putative surface HA protein harbored a typical Baculovirus gp64 envelope glycoprotein domain (Bac_gp64, e-value 4.64e^-48^), a signal peptide (SP) for transmembrane at the N terminus and a C terminal domain. We then surveyed the diverse libraries *D. gallinae* to assess the presence of these genome segments that were detected in all libraries following symmetrical expression patterns, based in FPKMs, supporting the possibility that the segments corresponded to the same virus. Expression levels were higher in fed deutonymph, lower in unfed protonymphs and interestingly, in each library, expression values of the structural segments were higher than the nonstructural ones; this was expected and is indirect evidence of active infection (**Table 2**). Some quaranjaviruses harbor additional segments, besides the core ones, associated with replication and infectivity. We used diverse strategies to identify more divergent segments based on levels of co-expression, relaxed e-value searches, or using the conserved terminal untranslated region as a signature tag of other segments. We did not find any evidence of additional segments. Based on these genomic data, genetic distance, functional and structural annotation, and expression profiles we suggest that these genome segments correspond to a novel virus, a putative member of a new species within the genus *Quaranjavirus* that we tentatively dubbed Red mite quaranjavirus 1 (RMQV1). To entertain this hypothesis, we generated evolutionary insights based on the phylogenetic analysis of PB1 and NP proteins of RMQV1 and diverse orthomyxoviruses. The resulting trees indicate that, clearly, RMQV1 clusters within the quaranjavirus clade, basally branching in an emergent subgroup of viruses recently found in metagenomic studies of ticks: Uumaja virus, detected in circumpolar seabird ticks, *Ixodes uriae* parasitizing penguins from northern Sweden ^96^, Zambezi tick virus 1 identified in *Rhipicephalus* sp. ticks from small ruminants collected in Mozambique ^97^ and Granville quaranjavirus, detected in *Amblyomma dissimile* ticks parasitizing hunted game animals (*Iguana iguana* and *Mazama americana*) from Trinidad and Tobago ^98^. There is no information regarding the biology and ecology of these viruses, nor any data that could suggest their potential veterinary or medical importance. Further work should address the potential impact of these viruses on their hosts and their ecological niche. In the specific case of RMQVY, this appears to be the first report of a quaranjavirus linked to mites, which raises more questions about its biology. It would be interesting to assess its prevalence, and whether this virus could be or is being transmitted to poultry.

**Figure 11.**
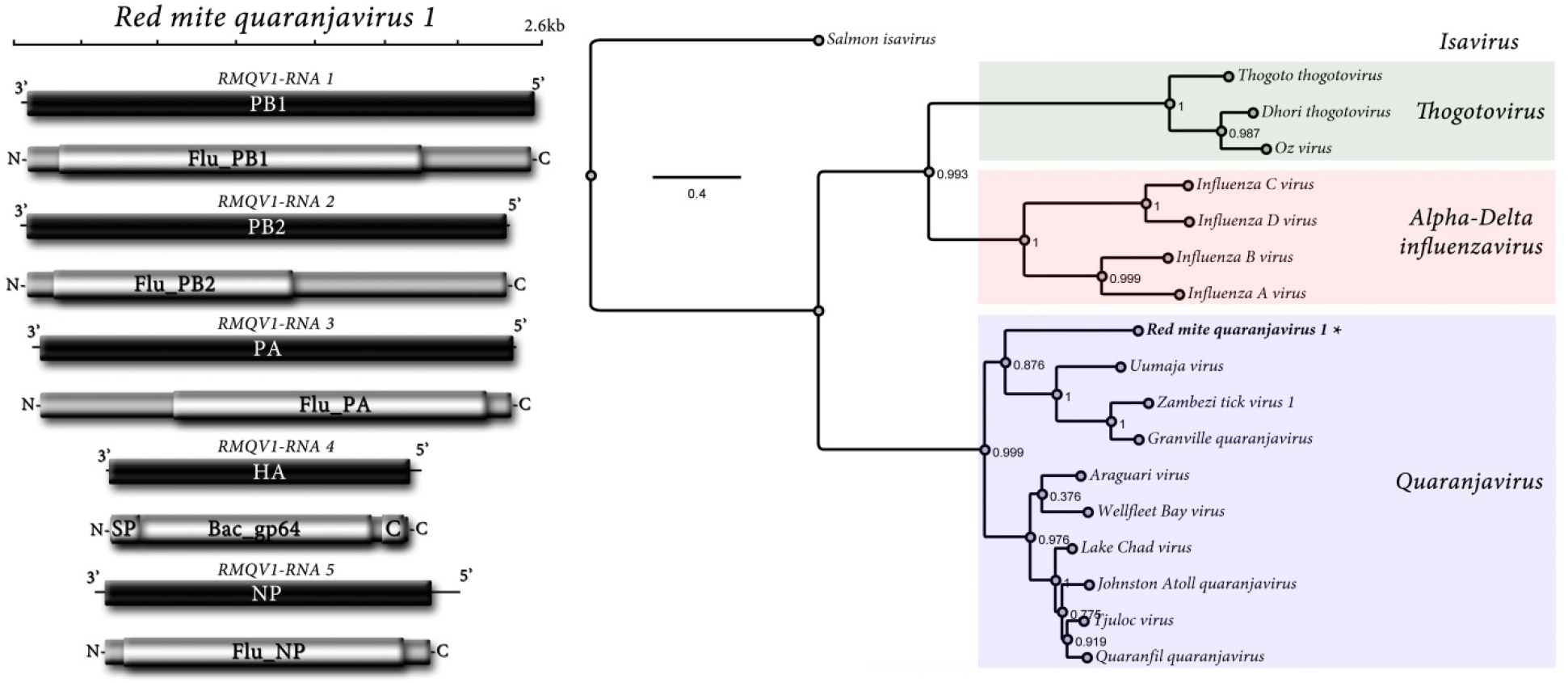
Red mite quaranjavirus 1. (Left panel) Genome graphs depicting genome segments and predicted gene products of Red mite quaranjavirus 1. Black rectangles indicate ORFs coding sequences, grey rectangles predicted proteins, light grey rectangles indicate coordinates of structural or functional domains. Abbreviations are described in the main text. (Right panel) Maximum likelihood phylogenetic tree based on alignment of predicted PB1 proteins of Red mite quaranjavirus 1 (asterisk) and related viruses. Branch labels represent FastTree support values. Scale bar represents substitutions per site.

For a more compehensive overview of the *D. gallinae* virome, please see Supplementary Data S17, with phylogenetic analyses of: red mite picorna-like virus, red mite densovirus, red mite virga-like virus, red mite associated cyclovirus, red mite associated hypovirus, and red mite associated cystovirus. The presence of viral sequences in libraries of unfed protonymphs (not exposed to host blood feeding) or dissected midguts (dissected interior tissue) demonstrate a genuine *D. gallinae* virome (**Table 3**), ruling out the possibility of host (blood/skin) or mite body surface environmental contaminants, respectively. In summary, a diverse and complex red mite virome was detected and characterized for *D. gallinae*, including novel viral species of several families of viruses, potentially having important impacts on the biology and health of their hosts. Future studies oriented to determine epidemiological, ecological and veterinary effects of these viruses are warranted.

**Table 3.** Expression values (FPKM) of viruses across libraries derived from individual developmental stages of red mites.

| *virus*<br>*taxa* | *libraries*<br>*-----*<br>*virus segments* | Unfed<br>Protonymph | Fed<br>Protonymph | Fed<br>Deutonymph | Fed<br>Adult | Midgut<br>(adult) |
| --- | --- | --- | --- | --- | --- | --- |
| *Densovirus* | *RMDEV1* | 12 | 544 | 1527 | 1768 | 1053 |
| *Picornavirus-like*<br>*virus* | *RMIV1* | 50485 | 25552 | 13748 | 15654 | 10258 |
|  | *RMDIV1* | 49261 | 53114 | 47501 | 23868 | 76529 |
| *Virga-like*<br>*virus* | *RMVLV1* | 2968 | 944 | 38301 | 60250 | 7069 |
|  | *RMVLV2* | 768 | 1987 | 12135 | 13266 | 15634 |
| *Cystovirus* | *RMACTV1-RNA1* | 39 | 18 | 21 | 14 | 16 |
|  | *RMACTV1-RNA2* | 571 | 416 | 299 | 498 | 326 |
| *Hypovirus* | *RMAHV1* | 165 | 75 | 10 | 46 | 19 |
| *Cyclovirus* | *RMACYV1* | 42442 | 138524 | 5456 | 1050 | 13279 |

## Conclusion

Despite *D. gallinae* being a global and highly debilitating poultry pest, the availability of high-throughput data and experimental platforms is lacking, with only rare exceptions ^24,99,100^. We have previously used Illumina RNA-seq data to describe transcriptomes of several tissues of the tick *I. ricinus* ^101-103^. *Here, we have sequenced and assembled new D. gallinae* developmental stage-specific transcriptomes, a key informative dataset enabling insights into molecular adaptions of this ectoparasite to its obligatory blood-feeding life-style. It also provides us with a catalogue of potential targets suitable for target-based acaricide development. The sequences of assembled contigs, annotations, and respective expression values are available through a user-friendly one-piece hyper-linked excel sheet (see Data availability). Unlike previous studies, our RNA-seq based work is complemented by viability assays, performed through the artificial membrane feeding platform and haemocoel microinjection, and also by mass spectrometry metabolite identification. Finally, we have identified the *D. gallinae*-specific RNA virome and have highlighted viruses of possible concern. This work presents an integrative assessment of assembled and curated Illumina-originating data, describing the molecular traits inherently linked to the blood feeding life style of *D. gallinae*, and reveals its emerging role as a reservoir of new viral variants.

## Materials and Methods

### Collection of mites, RNA extraction, and library preparation

Mites were collected by brushing cages of egg-producing hens in the International Poultry Testing Ustrasice (MTD Ustrasice, Czech Republic). Mites were briefly anaesthetised with CO_2_ using a FlyPad (Flystuff.com) and were separated into three developmental stages: protonymphs, deutonymphs, and adults. Protonymphs were further separated into unfed and blood-fed mites. Midguts were micro-dissected from adult stages. Mites were placed on a double-sticky tape, ventral side down, decapitated with a razor, and midguts were pulled out into a drop of DEBC-treated PBS. Whole mites of different developmental stages (each n = 30) were homogenised with an eppi pestle and midguts (n = 40) were homogenised using a 29G insulin syringe needle. Total RNA was extracted from the homogenates with a NucleoSpin RNA kit (Macherey Nagel) and eluted into RNase-free water, with a yield of 2 – 7 µg of total RNA. RNA was quality checked by Agilent 2100 BioAnalyser with RIN values ≥ 9. A non-stranded cDNA library was prepared by NEBNext® Ultra™ RNA Library Prep Kit for Illumina® and sequenced on NovaSeq600 by Novagene Co., Ltd. as 150 bp paired-end reads.

### Transcriptome assembly, data deposition, filtering of outputs

Transcriptome assembly and coding sequence extraction were carried out as described previously (Ribeiro et al., 2013). Briefly, reads were stripped of their contaminating primers, and bases with qual values < 20 were trimmed. Clean reads were assembled using the Abyss (Birol et al., 2009) and Trinity (Grabherr et al., 2011) assemblers. These assemblies were merged using a parallelised pipeline of blastn and cap3 assembler (Huang and Madan, 1999) as described previously (Karim et al., 2011). All open reading frames larger than 200 nucleotides were extracted and those matching known proteins or having a signal peptide were retained. The resulting peptide and coding sequences were mapped to a hyperlinked spreadsheet, including blastp and rpsblast matches to several databases, as well as an indication of the signal peptide (Nielsen et al., 1999), transmembrane domains (Sonnhammer et al., 1998) and O-galactosylation sites (Hansen et al., 1998). Transcripts assigned to a given developmental stage we filtered by 16 fold change factor, i.e. to denote a transcript “adult-specific”, its FPKM values need to >16× over its FPKM values in the transcriptome of fed protonymphs, and at the same time in the transcriptome of fed deutonymphs (Supplementary Data S1). To list transcripts enriched in the transcriptome of fed over unfed protonymphs, only transcripts with clear annotation (E value < e^-100^), coverage > 10%, and FPKM in fed protonymphs > 5 were considered. The transcripts were then listed according to highest fed / unfed ratios. If zero values were present in transcriptomes of unfed protonymphs, the transcripts were listed according to higher FPKM values in transcriptomes of fed protonymphs (Supplementary Data S2).

### Phylogenetic analyses

The dataset used for the phylogenetic analyses of arthropod vitellogenins consisted of 67 amino acid sequences representing homologues from ticks, mites, acarids, spiders, scorpions, horseshoe crabs, insects and crustaceans, with the latter being an outgroup (Supplementary Data S9). The vitellogenin sequences were retrieved as GenBank annotated entries or were mined from the genome/ transcriptome assemblies available in GenBank using the tBLASTn algorithm and E-value cutoff < 10^−5^. The structurally similar hemelipoglycoproteins of ticks (e.g., GenBank: ABK40086, ACF35055) and their related tick sequences annotated in GenBank as vitellogenins (e.g., GenBank: AXP34688, BAJ21514, AXP34687, XP_029826448) were not included in the analysis because these proteins are not true vitellogenin homologues. Vitellogenin sequences were aligned in MAFFT v7.017 implemented in Geneious Prime v2019.0.4 ^104^ using the automatic selection of the alignment strategy and default parameters for the gap opening penalty (1.53) and the offset value (0.123). Non-homologous regions of the alignment were trimmed so the final alignment comprised 3578 amino acid positions comprising the three structural domains typical for vitellogenins (i.e., the vitellogenin N-terminal region, domain of unknown function [DUF1943], and von Willebrand factor type D domain). The phylogenetic tree was reconstructed by the maximum likelihood (ML) method in IQ-TREE v1.6.12 ^105^ using the LG+F+R6 protein model selected by ModelFinder ^106^. Bootstraps were based on 1000 replicates. Bayesian inference (BI) analysis was performed in MrBayes v3.2.7a ^107^ implemented in CIPRES Science Gateway v3.3 ^108^ using four simultaneous MCMC chains sampled at intervals of 100 trees and posterior probabilities estimated from 2 million generations. The WAG+F+G4 model of evolution was selected by ModelFinder. The burn-in period represented 10% of all generations. The tree was visualized in Geneious Prime v2019.0.4 and graphically modified in Adobe Illustrator CS5.

The dataset used for the phylogenetic analysis of the cys-loop ligand-gated ion channel gene family consisted of 121 amino acid sequences that corresponded to the dataset of Dermauw et al. ^63^ and was enriched for the homologues from *D. gallinae* (Supplementary Data S10). The sequences retrieved from GenBank annotated entries were aligned and treated as described above for the vitellogenin dataset. Final alignment comprised 502 amino acid positions. The ML-based phylogenetic tree and its nodal supports were calculated using the LG+I+G4 protein model as described for the vitellogenins.

The dataset used for the phylogenetic analysis of the thioester-containing protein (TEP) family contained 29 amino acid sequences of *D. gallinae* TEPs and its invertebrate homologues from the fruit fly *D. melanogaster*, the hard tick *I. ricinus*, and the horseshoe crab *L. polyphemus* (Supplementary File S14). The sequences were aligned and treated as described above for the vitellogenin dataset. Final alignment of full amino acid sequences (∼1500 residues) comprised 2737 amino acid positions. The ML-based phylogenetic tree and its nodal supports were calculated using the BLOSUM62+G protein model as described for the vitellogenins.

### Metabolomic analysis of haem biosynthesis intermediates

For metabolomic analysis, mites were collected in a poultry house by brushing and stored in a 1 liter bottle with a paper lid at 21°C in the dark. Next day, a mixture of eggs, larvae, and unfed protonymphs was sorted out as described above. Then the sample of unfed stages (to prevent detection of heme intermediates of host origin), with a predominant fraction of unfed protonymphs, of 21.4 mg mites was extracted with 150 µl of a cold extraction medium MeOH:ACN:H_2_O (2:2:1 v/v/v) containing an internal standard 4-fluorophenylalanine (100 µl of 0.5 ng/µL). The sample was then homogenised using a Tissue Lyser II (Qiagen, Prague, Czech Republic) at 50 Hz, 0 °C for 5 min. The mixture was then centrifuged at 7000 RPM and 5 °C for 10 min, and the supernatant was filtered through a 0.2 μm PVDF mini-spin filter (HPST, Prague, Czech Republic) at 8000 RPM and 5 °C for 10 min. Finally, a 50 ul aliquot of the supernatant was used for liquid chromatographic – high resolution mass spectrometric analysis (LC-HRMS). The LC-HRMS methods were described in detail earlier^109,110^. Briefly: A Q Exactive Plus Orbitrap mass spectrometer coupled with a Dionex Ultimate 3000 liquid chromatograph (all Thermo Fisher Scientific, San Jose, CA, USA) was used for profiling of <u>haem biosynthesis intermediates</u>. Metabolites were separated on a 150mm × 4.6 mm i.d., 5μm, SeQuant ZIC-pHILIC (Merck KGaA, Darmstadt, Germany) with a mobile phase flow rate of 450 μL/min, an injection volume of 5 μL, and a column temperature of 35°C. The mobile phase was: A = acetonitrile, B = 20 mmol/L aqueous ammonium carbonate (pH = 9.2; adjusted with NH4OH); gradient: 0 min, 20% B; 20 min, 80% B; 20.1 min, 95% B; 23.3 min, 95% B; 23.4 min, 20% B; 30.0 min 20% B. The Q-Exactive settings were: Mass range 70-1000 Daltons; 70 000 resolution (m/z 200; 3 × 106 Automatic Gain Control (AGC) target and maximum ion injection time (IT) 100 ms; electrospray operated in positive mod: 3000 kV spray voltage, 350°C capillary temperature, sheath gas at 60 au, aux gas at 20 au, spare gas at 1 au, probe temperature 350°C and S-Lens level at 60 au. Data were processed using XcaliburTM software, version 4.0 (Thermo Fisher Scientific, San Jose, CA, USA). Compounds used in the study: Deionized water was prepared using a Direct Q 3UV purification system (Merck, Prague, Czech Republic). Methanol and acetonitrile (OptimaTM grade) were purchased from Fisher Scientific (Pardubice, Czech Republic); ammonium carbonate, 25% ammonia solution, 4-fluorophenylalanine, 5-aminolevulinate, haemin, porphobilinogen and protoporphyrin IX from Merck (Prague, Czech Republic).

### *Ex vivo* artificial membrane feeding

Mites were collected and stored as desribed above. After 14 days, vital individuals were selected and used for *ex vivo* feeding experiments. The feeding unit is a plastic cylinder (17 mm in diameter, 70 mm in length) horizontally divided in the middle by a membrane (silicone-impregnated Goldbeater’s skin) into two chambers. The upper chamber contains 2 ml of fresh, defibrinated (by mixing with 4 mm glass beads) chicken blood supplemented with tested inhibitor Torin2 (Sigma-Aldrich, catalogue number SML1224). The lower chamber of the membrane feeding device contained the mites. The feeding unit was placed in the dark in an incubator set at 41°C. After 5 hours, fed individuals were collected and stored in 1,5 ml microcentrifuge tubes (10 individuals in each) in an incubator set at 21°C. The mite’s viability and vitality was monitored every 24 hours for 7 days. Torin2 was dissolved in DMSO and further diluted to final concentrations: 500, 250, 100, 10, and 1uM. DMSO stocks were diluted in the blood meal to 1% v/v DMSO in the blood meals.

### Microinjection

Mites were collected as described above and freshly-fed adult females were sorted out. Mites were immobilised on an adhesive tape and a volume of 13,8 nl of tested compound (≤1% v/v DMSO) was injected into the mite’s hemocoel by microinjector (Drummond, USA). After 10 minutes, injected mites were collected into 1,5 ml microcentrifuge tubes (5 individuals in 5 eppi tubes for each concentration) and stored in the incubator set at 21°C. The mite’s viability and vitality was monitored every 24 hours for 7 days. Compounds used in the study: Fipronil (Sigma-Aldrich Supelco, catalogue number 16785), Fluralaner (Cayman Chemical, item number 22061), Ivermectin (Sigma-Aldrich, catalogue number I8898).

### Virus RNA identification and analyses

Virus discovery, detection and characterization was implemented as described elsewhere ^111^. In brief, transcriptome assemblies generated in this work were used as input for virus discovery. The complete NR release of viral protein sequences was retrieved from https://www.ncbi.nlm.nih.gov/protein/?term=txid10239[Organism:exp]. The integrated Red mite RNA assembly was assessed by multiple TBLASTN searches (max e-value = 1 × 10^−5^) using, as probe, the complete predicted non-redundant viral proteins in a local server. Significant hits were explored by hand, and redundant contigs were discarded. Potential virus like sequences were curated by iterative mapping of reads using Bowtie 2 v2.4.4 available at http://bowtie-bio.sourceforge.net/bowtie2/index.shtml. Open reading frames (ORF) were predicted by ORFfinder. Predicted proteins were subjected to BLASTP searches at NCBI and to domain-based Blast searches against the Conserved Domain Database (CDD) v3.19 implemented in https://www.ncbi.nlm.nih.gov/Structure/cdd/cdd.shtml and supplemented with SMART http://smart.embl-heidelberg.de/, Pfam http://pfam.xfam.org/ PROSITE http://prosite.expasy.org/ and HHPred https://toolkit.tuebingen.mpg.de/tools/hhpred to characterize more divergent functional domains. Signal and membrane peptides were assessed with SignalP v4.1 http://www.cbs.dtu.dk/services/SignalP/ Viral RNA levels were calculated with Cufflinks http://cole-trapnell-lab.github.io/cufflinks/ or alternatively with the Geneious suite 8.1.9 (Biomatters Inc.) as Fragments Per Kilobase of virus transcript per million mapped reads (FPKM). Evolutionary insights were resolved by MAFTT 7.310 alignments http://mafft.cbrc.jp/alignment/software/ of amino acid sequences of the predicted viral polymerases or capsid proteins which were used for phylogenetic analyses based on FastTree approximately maximum-likelihood phylogenetic trees http://www.microbesonline.org/fasttree/ with standard parameters. Support for individual nodes was assessed using an approximate likelihood ratio test with the Shimodaira-Hasegawa like procedure. Tree topology, support values and substitutions per site were based on 1,000 tree resamples. Most sequence analyses results were integrated and visualised using the Geneious suite 8.1.9 (Biomatters Inc.). Virus genome diagrams were designed using Adobe Photoshop C3 version 10.

## Data availability

The data are available as BioProject PRJNA597301, including nt and aa fasta files, and a hyperlinked excel sheet is available to download at https://proj-bip-prod-publicread.s3.amazonaws.com/transcriptome/Dermanyssus_gallinae/Derm_gallinae.zip. Viral sequences were deposited at the NCBI with the following GenBank accession numbers: RMACTV1-L,ON160022;RMACTV1-S,ON160023;RMQV1-HA,ON160024;RMQV1-NP,ON160025; RMQV1-PA,ON160026;RMQV1-PB1,ON160027;RMQV1-PB2,ON160028;RMDIV1,ON160029; RMIV1, ON160030;RMDEV1,ON160031;RMVLV1,ON160032;RMVLV2,ON160033;RMACYV1,ON160034;RMAHV1,ON 160035.

## Acknowledgement

We thank Martina Hajduskova (www.biographix.cz) for the graphical visualisation of Figure 2D.

